# Optimizing epitope conformational ensembles using *α*-synuclein cyclic peptide “glycindel” scaffolds: A customized immunogen method for generating oligomer-selective antibodies for Parkinson’s disease

**DOI:** 10.1101/2021.09.13.460126

**Authors:** Shawn C.C. Hsueh, Adekunle Aina, Andrei Yu. Roman, Neil R. Cashman, Xubiao Peng, Steven S. Plotkin

## Abstract

Effectively presenting epitopes on immunogens, in order to raise conformationally selective antibodies through active immunization, is a central problem in treating protein misfolding diseases, particularly neurodegenerative diseases such as Alzheimer’s disease or Parkinson’s disease. We seek to selectively target conformations enriched in toxic, oligomeric propagating species while sparing the healthy forms of the protein that are often more abundant. To this end, we computationally modelled scaffolded epitopes in cyclic peptides by inserting/deleting a variable number of flanking glycines (“glycindels”), to best mimic a misfolding-specific conformation of an epitope of *α*-synuclein enriched in the oligomer ensemble, as characterized by a region most readily disordered and solvent-exposed in a stressed, partially denatured protofibril. We screen and rank the cyclic peptide scaffolds of *α*-synuclein *in silico* based on their ensemble overlap properties with the fibril, oligomer-model, and isolated monomer ensembles. We present experimental data of seeded aggregation that supports nucleation rates consistent with computationally predicted cyclic peptide conformational similarity. We also introduce a method for screening against structured off-pathway targets in the human proteome, by selecting scaffolds with minimal conformational similarity between their epitope and the same solvent-exposed primary sequence in structured human proteins. Different cyclic peptide scaffolds with variable numbers of glycines are predicted computationally to have markedly different conformational ensembles. Ensemble comparison and overlap was quantified by the Jensen-Shannon Divergence, and a new measure introduced here—the embedding depth, which determines the extent to which a given ensemble is subsumed by another ensemble, and which may be a more useful measure in developing immunogens that confer conformational-selectivity to an antibody.

**Graphical TOC Entry:** 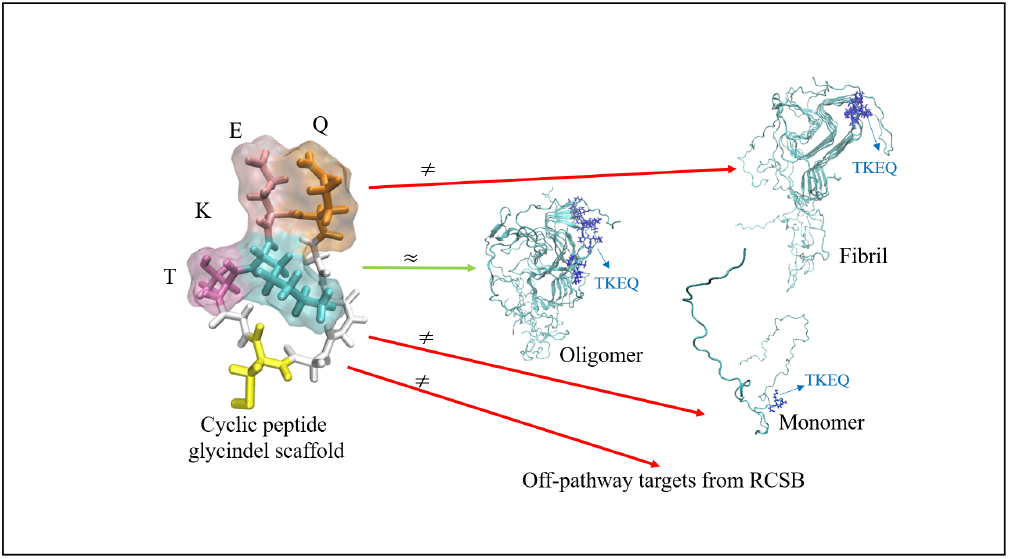

## 1 Introduction

A key step in the development of a therapeutic antibody or active vaccine is the immunization strategy (*1*), namely, the choice of protein epitope and how it will be presented to an animal or human immune system. Both primary sequence and conformation of the epitope determine the particular protein morphologies to which the resulting antibodies will be selective.

Nowhere has conformational-selectivity been more important to immunotherapies than in protein-misfolding diseases (*2*). For this class of diseases, an effective antibody must be able to spare healthy protein and discriminate misfolded protein species that lead to molecular and cellular pathology (*1*). Since the primary sequences of healthy and aberrant protein are generally the same, barring splice variants and perhaps post-translational modifications, the efficacy of an antibody is then due to its selective preference for binding to a misfolded conformation over healthy in-vivo “native” conformations.

For many proteins involved in misfolding disease however, the native conformational ensemble is intrinsically disordered (*3*). Examples include amyloid-*β* (A*β*), tau protein, *α*-synuclein, and the low-complexity domains in FUS and TDP43. Raising an antibody that avoids the majority of diverse conformations of an intrinsically-disordered protein’s ensemble is a challenge. For example, a peptide consisting of a contiguous fragment of native primary sequence tethered to an immunogen such as keyhole limpet hemocyanin (KLH) will likely exhibit overlap in its presented ensemble with the ensemble of isolated native monomer.

Many neurodegenerative diseases, including Alzheimer’s disease (AD), Parkinson’s disease (PD), chronic traumatic encephalopathy (CTE) and amyotropic lateral sclerosis (ALS), spread throughout the brain via a prion-like mechanism involving soluble oligomers (*1, 4* – *11*) Soluble oligomers contain roughly 4-40 chains of protein, which exist in a misfolded conformational ensemble that is conformationally labile and very difficult to experimentally characterize. We computationally model some key aspects of the oligomer ensemble using molecular dynamics, in order to predict disease-specific epitopes.

Cyclic peptides, a subclass of macrocycles, are polymers of amino acids that have been conjugated to form a ring-like topology (Fig. 1). They have been increasingly used as therapeutics, often as small molecule drugs to bind targets (*12* –*16*). They have also been used as immunogens (*17* –*22*) to raise oligomer-selective antibodies in Alzheimer’s disease using the method that we describe here (*19, 23, 24*).

**Figure 1:**
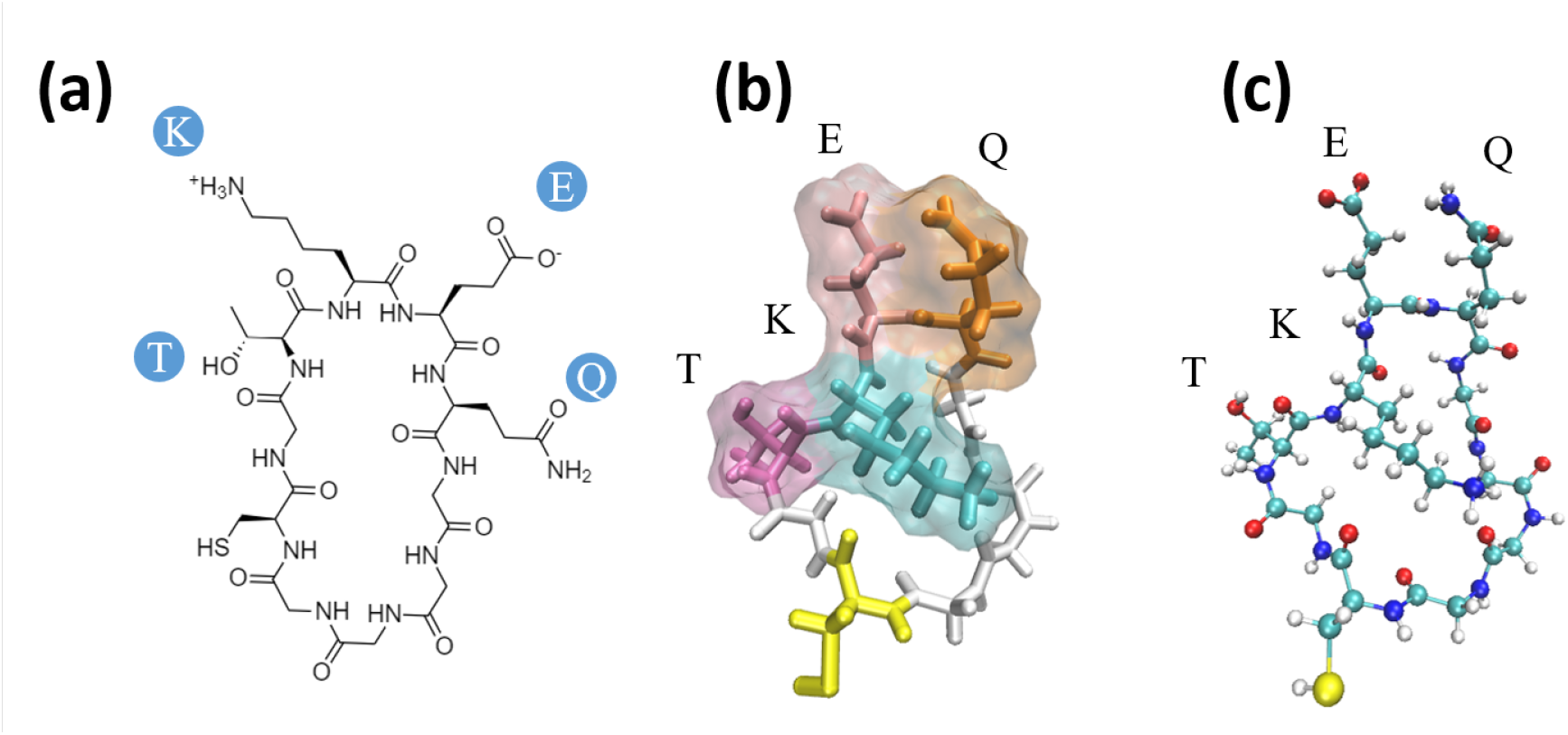
Cyclic peptide renderings for cyclo(CGTKEQGGGG), a scaffold of TKEQ. (**a**) 2D representation of the cyclic peptide. (**b**) 3-dimensional rendering of the cyclic peptide in licorice, also showing the surface of the TKEQ epitope. Colors are assigned by residue name, with glycine in white, cysteine in yellow, threonine in dark pink, lysine in cyan, glutamate in light pink, and glutamic acid in orange. (**c**) Ball and Stick (CPK) rendering with color assigned by the atom identity.

In this paper, we do not address the non-trivial problem of misfolding-specific epitope prediction, which we have treated elsewhere (*25*) (see Methods Section 3.1). We focus instead on the problem of how to properly present a predicted epitope to the immune system of an animal (or human, in the case of active immunization), so that the resulting antibodies generated by the animal are selective to disease-specific forms of the protein.

We start from computationally generated ensembles of the *α*-synuclein fibril, isolated monomer, and stressed, partially disrupted proto-fibril as defined below, which is used as a model for the toxic oligomer—a species we wish to target with conformationally selective antibodies. The *α*-synuclein oligomer model was generated computationally. Although oligomer structures involving select portions of the primary sequence constrained into peptide macrocycles have been crystallized (*26*), no full-length *α*-synuclein oligomer structure or ensemble, or partial length *α*-synuclein oligomer structure containing our epitopes of interest, has been experimentally characterized at this time. We briefly review the epitope prediction method described in (*25*), which is applied here to an *α*-synuclein protofibril. Given an epitope and an oligomer model, we then construct various cyclic peptide immunogen constructs of that epitope by varying the number of glycines flanking the epitope on the N- and Ctermini, which we term “glycindel” scaffolds. Here we introduce only glycines as extrinsic amino acids, as they are relatively non-immunogenic, which avoids potential immunological targeting of regions outside of the epitope of interest. Within this restricted space of extrinsic sequence and structure, we determine the optimal scaffolding of the cyclic peptide construct, by maximizing the overlap with the stressed fibril ensemble and minimizing the ensemble overlap with either the monomer or the unstressed “native” fibril. We also minimize the tendency of antibodies to a scaffolded epitope to elicit off-pathway targeting, by minimizing the conformational similarity to unrelated structured proteins in the human proteome that contain the epitope’s primary sequence. Using the above features as screening criteria, we used a ranking method we have developed previously (*27*) involving multi-criteria decision making analysis to systematically rank the scaffolds from best to worst. Such a ranking allows for the experimentalist to construct a reduced number of cyclic peptide scaffolds that are most likely to have the desired features of eliciting antibodies with conformationally selective binding to toxic oligomer species.

The pipeline of the *in silico* screening method is given in Fig. 9. Each step is described in the Methods. In the Results, we visually depict the projection of simulated ensembles in reduced conformational space, and we formalize the ensemble overlap. We then develop the above-described method to find off-pathway targets for a given scaffold, and rank 48 candidate cyclic peptide scaffolds. We show that the same similarity metrics used in the glycindel ranking can also explain the seeding activity of cyclic peptides. We finally discuss the non-trivial question of weight assignments for ranking candidates, alternate scaffolding methods, the validity of the virtual screening method, the benefits of *in silico* screening to facilitate more efficient *in vitro* and *in vivo* screening, and the sensitivity or robustness of epitope prediction depending on the structural model of the fibril used in the epitope prediction algorithm.

## 2 Results and Discussion

Using our previously developed epitope prediction algorithm (*25*), amino acid sequence EKTKEQ (residues 57-62 in *α*-synuclein) was predicted as a misfolding-specific epitope (*28*) (Fig. 10). To dissect the key amino acids in this epitope, we have analyzed three separate contiguous sequences subsumed by the epitope, namely EKTK, KTKE and TKEQ.

### 2.1 Comparing ensembles in reduced conformational space

In order to determine epitope scaffolds that may be capable of eliciting antibodies that are conformationally selective to soluble oligomers, the conformational ensemble of an epitope in a cyclic peptide scaffold is compared with three other ensembles of the epitope, namely, the monomer, fibril, and the stressed (partially disordered) fibril. The stressed fibril is taken as a proxy for the conformational ensemble of the epitope in a soluble oligomer, as successfully demonstrated previously (*23, 24*). A desired cyclic peptide scaffolded epitope construct that is oligomer-selective would have high ensemble similarity to the stressed fibril, and as well, low ensemble similarity to the equilibrium fibril and isolated monomer.

To determine the similarity between the four epitope ensembles (i.e. scaffold, stressed fibril, fibril, and monomer), we first generated these ensembles by performing molecular dynamics (MD) simulations (see Methods Section 3.3). The conformational similarity of the epitope between ensembles is quantified in a reduced conformational space of the epitopes (see Methods Section 3.4). For example, Fig. 2 shows the structural distributions in 1D for the four ensembles of epitope TKEQ, in two scaffolds. The two scaffolds chosen for this illustration are cyclo(CGTKEQGGGG) or (1,4)TKEQ, which stands for 1 glycine N-terminal to the TKEQ epitope and 4 glycines C-terminal to the epitope, and cyclo(CGGTKEQGGG), or (2,3)TKEQ.

**Figure 2:**
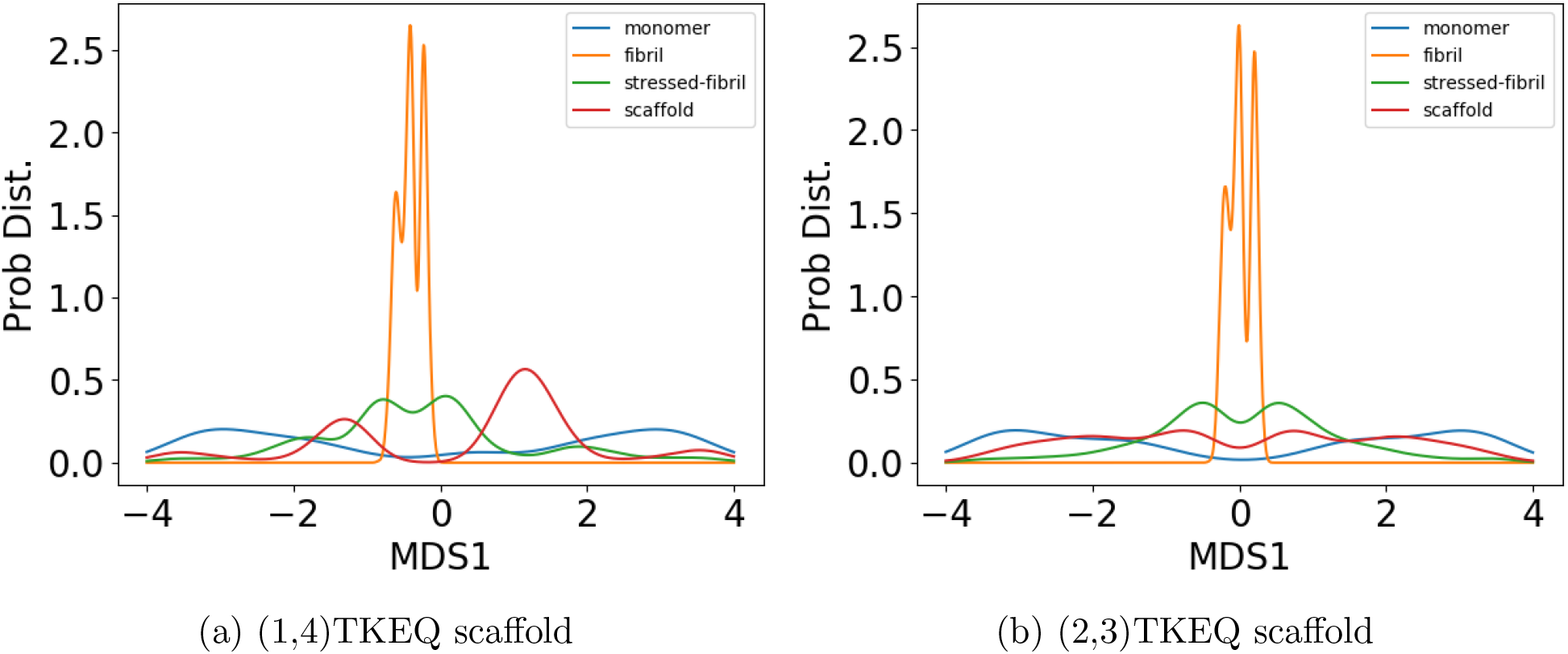
The equilibrium ensemble distributions for the TKEQ epitope projected by the multidimensional scaling (MDS) method (*32*) onto the first MDS dimension. For a given epitope, different cyclic peptide scaffolds possess different distributions, which will result in different overlap with the other three ensembles. By comparing the degree of ensemble overlap, the conformational selectivity of a scaffold can be assessed. The scaffolds shown are (1,4)TKEQ in panel (a), and (2,3)TKEQ in panel (b).

Although information may be lost when projecting high dimensional ensembles to lower dimension, the distributions in Fig. 2 do illustrate how the ensembles overlap. The fibril ensemble consists of a predominant sharp spike because of its rigid structure. On the other hand, the monomer ensemble is broadly distributed because it is natively unstructured and conformationally diverse. The stressed fibril ensemble distributes around the fibril ensemble because it is generated by forcing the partial unfolding of fibril by biased MD. Each cyclic peptide scaffold possesses a different distribution. As well, the stressed fibril, monomer, and fibril ensembles are slightly different in each case of Fig. 2, because all structural distributions are distance distributions based on RMSD, which are different for each scaffold ensemble. From the degree of overlap or similarity between a scaffolded cyclic peptide ensemble and the other ensembles, we can assess whether the scaffold has the potential to raise oligomer-selective antibodies while sparing fibril and monomer.

For example, both (1,4)TKEQ and (2,3)TKEQ scaffold ensembles have very low overlap with the fibril ensemble (3% and 8% respectively) (see Methods Section 3.5). Also, they both have high overlap with the stressed fibril, where (2,3)TKEQ is somewhat higher than (1,4)TKEQ (68% and 47%). The above (low fibril overlap, high stressed fibril overlap) are desired properties of a conformationally selective immunogen. On the other hand, the scaffolds also have high overlap with isolated monomer ensemble, with (2,3)TKEQ having much higher overlap than that in (1,4)TKEQ (74% and 47%). High overlap with the monomer ensemble is not a favorable property because of the possibility of targeting healthy protein. Naïvely from the 1D overlap measure, (1,4)TKEQ may be better than (2,3)TKEQ, since it has a higher ratio of stressed fibril ensemble overlap to monomer ensemble overlap (1.0 for (1,4)TKEQ *vs*. 0.92 for (2,3)TKEQ). However, with a more rigorous similarity measure analysis allowing for higher projected dimensions, and the ranking algorithm in Section 3.8, (2,3)TKEQ actually ranks higher than (1,4)TKEQ.

The similarity or overlap between ensembles is rigorously quantified by two measures, Jensen-Shannon Divergence (JSD) and embedding depth (see Methods Section 3.6). JSD is a measure that represents the dissimilarity between two ensembles, which has been used previously to compare protein ensembles (*29, 30*). Embedding depth represents the extent to which a given ensemble is subsumed by another, i.e. to what extent conformations in one ensemble are contained within another ensemble. It is a non-reciprocal measure that is introduced in Methods Section 3.6.1 to compare protein conformational ensembles (see also Ref. (*31*)). For example, the embedding depth of the fibril ensemble within the stressed fibril ensemble in Fig. 2a is 0.338, because fibril conformations are contained within the stressed fibril ensemble. Note that embedding depth between two identical ensembles is 0.5, so 0.338 has represented a large degree of embedding. On the other hand, the JSD between the two ensembles is 0.984, which represents two almost completely dissimilar ensembles. The efficacy of a scaffold to target a conformational species may thus likely be better represented by the embedding depth than the JSD.

### 2.2 Scaffold screening criteria and scaffold ranking

In Section 2.1, we showed that the various similarity measures that we had defined between a scaffolded cyclic peptide epitope and other ensembles can be optimized within the glycindel sequence space by varying the number of flanking N-terminal and C-terminal glycines (See Methods Section 3.2). The ensemble overlap of a given epitope with other ensembles can be tuned by changing the scaffolding residues. We thus computationally constructed 4 *×* 4 = 16 scaffolds for each epitope, corresponding to the epitope being flanked by anywhere between 1 and 4 glycines on each terminus or “side”, or 16 *×* 3 = 48 total for epitopes EKTK, KTKE, and TKEQ.

Fig. 3 shows the results for three *α*-synuclein epitopes; EKTK, KTKE and TKEQ, illustrating changes that can occur in ensemble similarity across all 48 scaffolds, as measured by Jensen-Shannon Divergence JSD (see Methods Section 3.6.2), embedding depth 𝒟_*A*|*B*_ (see Methods Section 3.6.1) and off-pathway target criteria OP (see Methods Section 3.7), by varying the number of flanking N-terminal and C-terminal glycines. Detailed values are in Table S1. The changes in the values corresponding to the fibril ensemble (JSD_cyclic-fibril_, and 𝒟_cyclic-in-fibril_) are all very small on the scale of the plots, so they appear to remain unchanged across all scaffolds. In practice, this means that these two criteria do not contribute significantly to the ranking between scaffolds. For epitopes EKTK and KTKE, there is a generally decreasing trend of JSD with increasing scaffold length (total residue number in the cyclic peptide), and a generally increasing trend for 𝒟 with increasing scaffold length. These trends are less significant for epitope TKEQ. We address this phenomenon further in Section 2.4.

**Figure 3:**
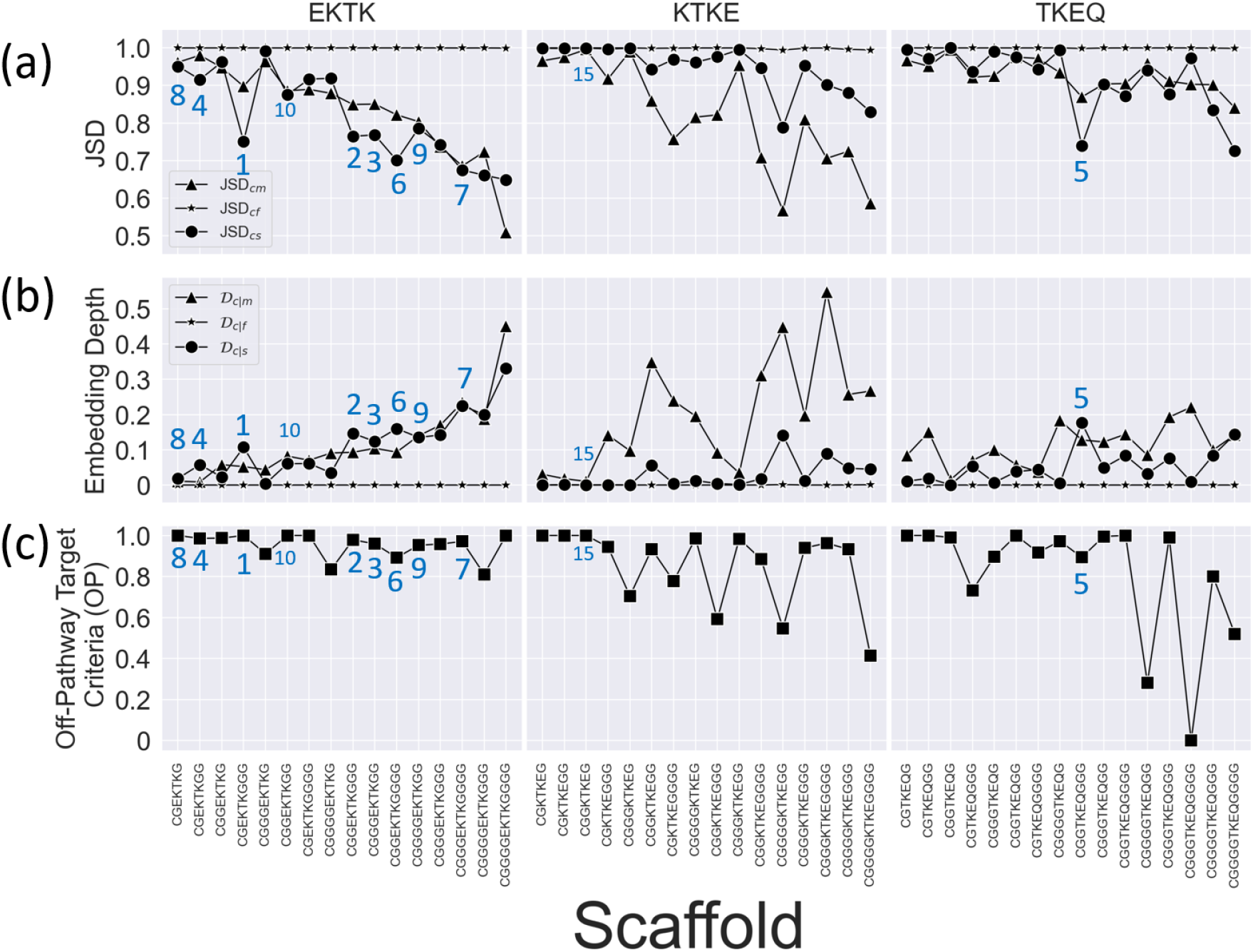
Measures for the rankings of all 16 epitope scaffolds, for three overlapping 4 residue sub-epitopes of EKTKEQ in *α*-synuclein (top of each column). (a) Scaffolded cyclic peptide ensemble dissimilarity to monomer (triangle), fibril (star), and stressed fibril (circle) ensembles, as measured by Jensen-Shannon Divergence (JSD), showing the changes in ensemble overlap with varying numbers of flanking glycines. (b) Scaffolded cyclic peptide ensemble embedding depth within the monomer (triangle), fibril (star), and stressed fibril (circle) ensembles, showing the changes in ensemble embedding with varying number of flanking glycines. (c) Normalized off-pathway targeting values (OP) for scaffolds with varying number of flanking glycines. Higher values indicate there is less predicted off-pathway targeting by a given scaffold. The ranks of the top 10 scaffolds are indicated in the figure panels, along with the rank for the highest ranking scaffold (15) for epitope KTKE.

The difficulty in manually assessing good scaffolds from multiple similarity measures led us to apply a systematic ranking method for the selection of the best performing scaffolds. The performance of each scaffold is assessed by 7 ranking criteria, including JSD_cyclic-fibril_, JSD_cyclic-monomer_, 1-JSD_cyclic-stress_, 1-𝒟_cyclic-in-fibril_, 1-𝒟_cyclic-in-monomer_, 𝒟_cyclic-in-stress_ and OP (the off-pathway target criterion). We formulate the ranking such that large values in all criteria are desired, so we thus subtract some JSD values from 1 is to convert dissimilarity to similarity, and we subtract some and 𝒟 values from 1 to convert similarity to dissimilarity. Scaffolds are ranked using the SMAA-TOPSIS algorithm (*27*) using these 7 criteria (see Methods Section 3.8). In the ranking algorithm, each criterion is assigned a weight for its relative importance; The weights used here are given in Table 1. Since JSD_cyclic-monomer_ and 1-𝒟_cyclic-in-monomer_ both represent the effect of monomer dissimilarity, the importance of monomer in the ranking amounts effectively to the sum of the two weights. The same applies to 1-JSD_cyclic-stress_ and 𝒟_cyclic-in-stress_, as well as JSD_cyclic-fibril_ and 1-𝒟_cyclic-in-fibril_. Thus, the importance respectively for fibril, monomer, stressed fibril, and off-pathway target criteria are 2, 2, 1, 1.5. The ranking for all 48 scaffolds is shown in Table S1. Such a ranking provides a therapeutic development strategy to predict which cyclic peptide scaffolds may be most promising to use in an active immunization for antibody generation to oligomer targets of *α*-synuclein.

**Table 1:**
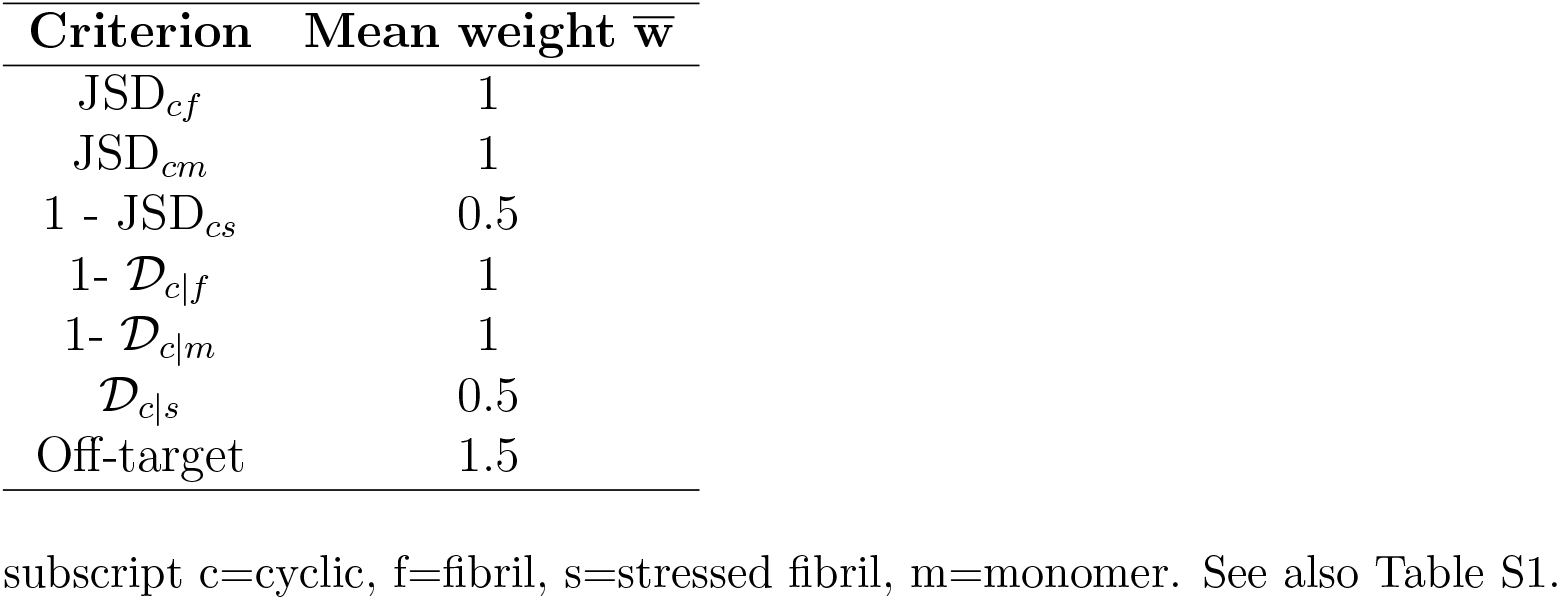
Mean weights of ranking criteria.

### 2.3 Embedding depth as a similarity measure compared with JSD

The various ensemble similarity measures are compared with each other in Fig. 4, which shows matrices of the Pearson’s correlation coefficient values, across all the scaffolds in

**Figure 4:**
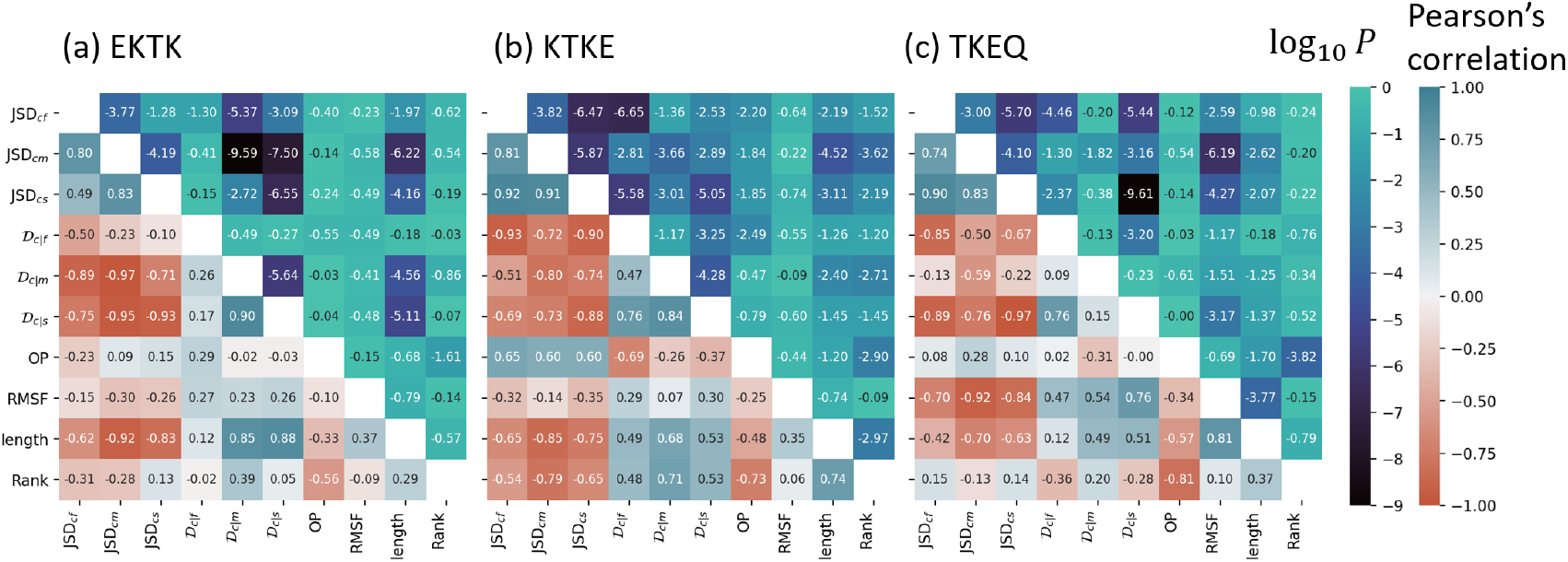
Correlation matrices of the ensemble comparison metrics JSD and 𝒟, and three other scaffold properties: Dynamic flexibility (RMSF) of an epitope, total residue length of the cyclic peptide scaffold, and the ranking, for *α*-synuclein epitopes (a) EKTK, (b) KTKE, and (c) TKEQ. The number in each square is the Pearson correlation coefficient.

Table S1. JSD and embedding depth 𝒟 are compared for all scaffolds, along with three other scaffold properties: Root Mean Squared Fluctuation of the epitope (RMSF), scaffold total residue length, and the ranking of each scaffold. JSD is calculated between ensembles by weight averaging values from 3 to 11 dimensions as described in Section 3.6.2; Embedding depth 𝒟 is calculated in 3-dimensions (3D) as described in Section 3.6.1. The three JSDs (JSD_cyclic-stress_, JSD_cyclic-fibril_, JSD_cyclic-monomer_) have mutually positive correlations, as shown in the top-left 3 *×* 3 matrices in Fig. 4a, b and c. This indicates that these JSD values may have implicit dependencies on one another and may contain some degree of redundant information. For epitope TKEQ, the JSD correlations between cyclic peptide ensembles and fibril, stressed fibril, and monomer ensembles may result at least in part from their flexibility, i.e. more flexibility of a cyclic peptide allows more conformational diversity, allowing in turn for more similar conformations to the other ensembles, reducing the JSD. Conversely, a rigid scaffold would have a narrow structural distribution, and thus minimal overlap with any of the other ensembles, resulting in larger JSD. The JSD values of various scaffolds thus have strong negative correlation with the RMSF for this epitope (8th column/row in Fig. 4c). This anticorrelation of RMSF with JSD is recapitulated partially by the (weaker) correlation of RMSF with 𝒟. This effect may indicate a systematic deviation of conformational similarity for scaffolds of this epitope, and may suggest better targeting can be achieved for EKTK and KTKE, where the anti-correlation of JSD with RMSF is less significant.

Embedding depth can help elucidate the implicit effects of scaffold rigidity. A rigid scaffold that is deeply embedded within the stressed fibril would be a desirable candidate. Compared across scaffolds for a given epitope as in Fig. 4, an anticorrelation between 𝒟_*c*|*s*_ and RMSF would be ideal for oligomer targeting. This is not observed for any of our epitopes, but the correlations are not significant for EKTK and KTKE, (*p* = 0.33 and *p* = 0.25) while for TKEQ the significance of the correlation of 0.76 is *p* = 0.0007. It is also worth noting that we are integrating a significant amount of information when investigating correlations across all scaffolds for a given epitope, in order to compare one epitope with another. The variance scaffold to scaffold for a given epitope is sufficiently large to allow for highly ranked scaffolds for any of the epitopes. That said, the median rankings for EKTK, KTKE, and TKEQ are 10.5, 37.5, and 25 respectively, and 9 of the top 10 scaffolds are for EKTK.

### 2.4 Scaffold properties and their impact on performance

The top 5-ranked scaffolds—(1,3)EKTK, (2,3)EKTK, (3,2)EKTK, (1,2)EKTK, and (2,3)TKEQ— all have substantially lower JSD_cyclic-stress_ than JSD_cyclic-monomer_, higher 𝒟_cyclic-in-stress_ than 𝒟_cyclic-in-monomer_, and high off-pathway (OP) target values (less off-pathway target severity) (Fig. 3 and Table S1). JSD_cyclic-fibril_ is approximately 1 for all scaffolds, and likewise 𝒟_cyclic-in-fibril_ ≈ 0, so these screening criteria cannot discriminate scaffolds.

We found that scaffolds with more residues tend to have smaller JSDs, and larger embedding depths 𝒟 (Fig. 3 and Fig. 4 “length” row/column). The trend is particularly apparent for EKTK scaffolds. It is likely that this trend is due to scaffold rigidity, since larger cyclic peptides have less structural constraint and more flexibility, allowing them to have higher overlap with the other disordered ensembles. On the other hand, while RMSF correlated positively with peptide length for all epitopes, only TKEQ achieved statistical significance (Fig. 5a), suggesting that the primary sequence of the epitope is as important as scaffold size in determining its dynamics and flexibility, and thus its potential similarity to other ensembles. As another indicator that primary sequence determines epitope properties, scaffold size itself does not correlate significantly with a scaffold’s ranking performance for TKEQ and EKTK (Fig. 5b), primarily because it is the *relative* overlap with the stressed fibril and monomer ensembles that determines the ranking. For KTKE, the correlation of scaffold size with ranking is such that smaller scaffolds tend to perform significantly better (Fig. 5b), mostly because the strongest trend with increasing scaffold length is increased overlap with the monomer ensemble (Fig. 3), which is an undesirable trait. The rankings themselves for KTKE are generally lower than those for the other two epitopes.

**Figure 5:**
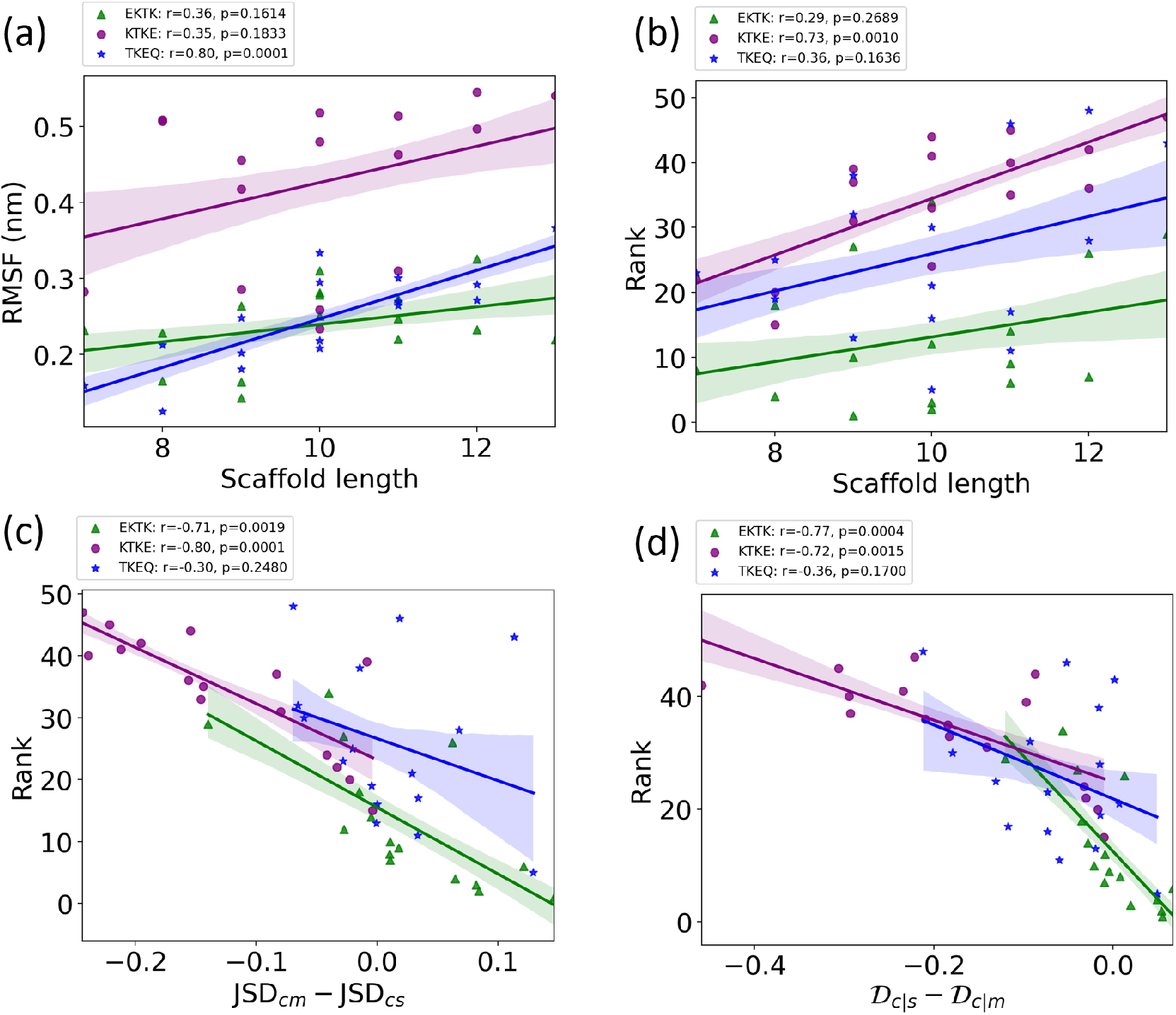
Epitope-dependent correlation between (a) RMSF and scaffold length (number of residues in the scaffold), (b) ranking and scaffold length, (c) rank and the quantity JSD_*cm*_ JSD_*cs*_, and (d) rank and 𝒟 _*c*|*s*_ *−*𝒟_*c*|*m*_, for *α*-synuclein epitope scaffolds. The Pearson correlation coefficient, *r* and the corresponding *p*-values are given for EKTK (green triangles), KTKE (purple circles) and TKEQ (blue stars) scaffolds. The shaded areas around the fitted lines are the 68% confidence intervals corresponding to the standard errors.

Plotting the scaffold rank *vs*. the *difference* in the JSD_*cm*_ *−*JSD_*cs*_, as well as the difference 𝒟_*c*|*s*_*−*𝒟_*c*|*m*_ (Fig. 5c,d) gives an indication of the importance of the relative overlap between the stressed-fibril *vs*. the monomer ensembles. Both abscissae in Fig. 5c,d are plotted as desirable quantities—the larger the value, the better the performance (higher rank or lower numerical rank value). For epitopes EKTK and KTKE, there is a significant correlation between a scaffold’s rank and these relative overlap measures. For epitope TKEQ, the correlation is not significant however, essentially because the variation in the off-pathway (OP) target criterion strongly affects the ranking for this epitope. This can be seen by retaining only those TKEQ scaffolds with *OP* above a threshold such as 0.85 for example, and measuring the correlation for this reduced, filtered dataset. This reduced dataset consists of 11 out of 16 scaffolds and yields *r* = *−*0.68, *p* = 0.019 in Fig. 5c and *r* = *−*0.88, *p* = 0.0002 in Fig. 5d.

Some epitopes tend to have better performance than others. EKTK and TKEQ scaffolds are generally ranked higher than KTKE scaffolds. As mentioned above, the median rankings for EKTK, TKEQ and KTKE are 10.5, 25, and 37.5 respectively. For the top 20 scaffolds, there are 12 EKTK scaffolds, 6 TKEQ scaffolds, and only 2 KTKE scaffolds, and 9 of the top 10 scaffolds are for EKTK, with only one TKEQ scaffold ranked 5th (Fig. 3). The poor performance of KTKE scaffolds appears to be due to their systematically lower structural similarity to the stressed fibril than to the unstructured monomer ensemble, as shown in Fig. 3. The higher monomer similarity may itself be due to the higher conformational flexibility (RMSF) of KTKE scaffolds, which is roughly twice as high on average as the RMSF of epitopes EKTK or TKEQ (Fig. 5a). I.e. flexibility in the scaffold construction could favor greater similarity to the monomer ensemble, because the monomer ensemble is inherently more conformationally diverse than that of the stressed fibril.

Since a cyclic peptide has an inherent curvature which may more strongly resemble the conformations of an epitope in a turn or bend, the averaged virtual bond angles representing the local curvatures of epitopes in the stressed fibril ensemble are calculated and compared. The virtual bond angle is defined as 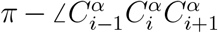 in the kink model (*33* –*35*), where 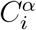 is the *C*_*α*_ atom of the *i*th amino acid. This angle has been shown to represent curvature in the continuum limit (*36*). However, the curvature of KTKE (1.13 *±* 0.14) is not significantly lower than that of EKTK (1.18 *±* 0.18) or TKEQ (1.10 *±* 0.16). As a result, the curvature itself does not explain the lower performance of KTKE.

### 2.5 Finding potential off-pathway targets

It is desirable to avoid off-pathway targets to minimize unwanted side-effects. To find the prevalence of unwanted targets in the proteome for a given epitope, we search through the RCSB database of resolved protein structures, to find proteins that might be potential off-pathway targets of antibodies that could be generated by each cyclic peptide scaffold (see Methods Section 3.7).

To illustrate the procedure, we demonstrate the off-pathway target analysis of (1,4)TKEQ scaffold here. Fig. 6a shows the scaffold ensemble for (1,4)TKEQ, projected on the first MDS coordinate, along with all entries of the human proteome having known PDB strutures and containing TKEQ motifs. The degree a PDB entry is embedded in the cyclic peptide ensemble is quantified as the embedding depth 𝒟_off-target-in-cyclic_, and is recorded as a percentage. Note that Fig. 6a is only for visualization; The formal calculation of embedding depth, 𝒟_off-target-in-cyclic_, is performed in 5 dimensions (5D). Nevertheless, we can see from this figure that the epitope in the context of the various PDB structures is conformationally distinct from the epitope in the context of the cyclic peptide.

**Figure 6:**
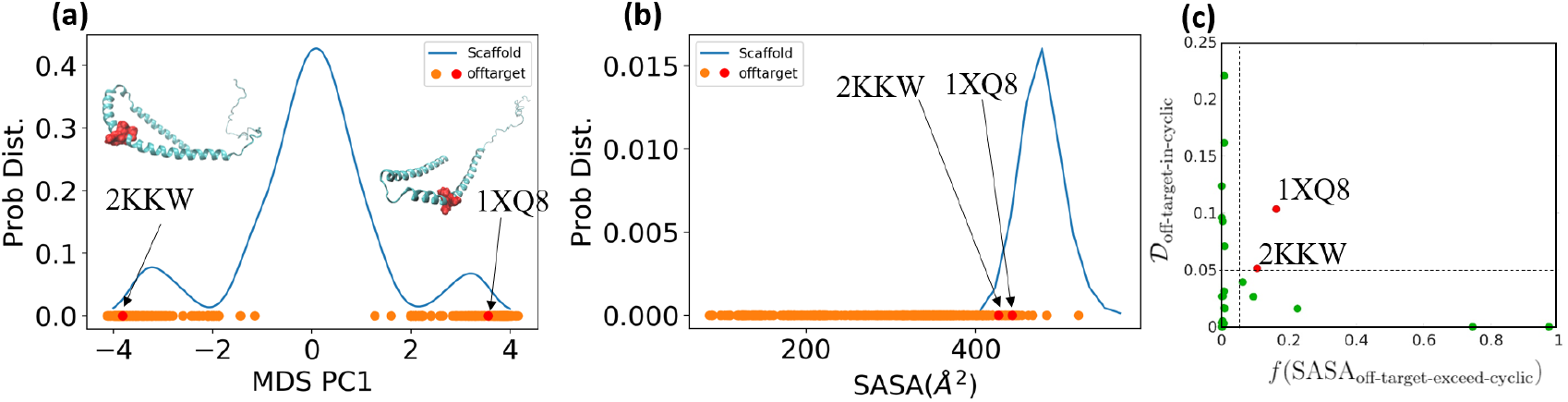
Off-pathway target analysis for (1,4)TKEQ. (a) Structural ensemble distribution of cyclic peptide (1,4)TKEQ in 1D along the first MDS component of the ensemble, along with the projected embedding of potential off-pathway targets. Most of the off-pathway targets are located at the periphery of the scaffold distribution. The actual calculation is performed in 5D. Structures of PDB entries 2KKW and 1XQ8 are rendered in ribbon schematics, and the epitope is rendered in red Van der Waals surface. (b) The SASA distribution of (1,4)TKEQ, along with the SASA for the off-pathway target structures. (c) Only 1XQ8 and 2KKW show both noticeable structural similarity (𝒟_off-target-in-cyclic_(5𝒟) *>* 5%) and SASA exposure (*f* (SASA_off-target-exceed-cyclic_) *>* 5%)

Fig. 6b indicates the distribution of the SASA for the (1,4)TKEQ ensemble, along with the SASA of TKEQ for all PDB entries in the human proteome with TKEQ motifs. The SASA of the scaffolded cyclic peptide ensemble ranges from 400Å^2^ to 600Å^2^, and serves as a good reference from which to compare. The fraction of the cyclic peptide ensemble that has lower SASA than a given PDB entry is defined as *f* (SASA_off-target-exceed-cyclic_).

We apply cutoff thresholds of 5% for both 𝒟_off-target-in-cyclic_ and *f* (SASA_off-target-exceed-cyclic_) to all PDB entries containing the epitope. In the case of (1,4)TKEQ, two PDB entries (1XQ8 (*37*) and 2KKW (*38*)) show *both* noticeable structural similarity (𝒟_off-target-in-cyclic_ *>* 5%) and solvent exposure (*f* (SASA_off-target-exceed-cyclic_) *>* 5%) relative to the (1,4)TKEQ ensemble (Fig. 6c and (1,4) entry in Fig. S2c). Other off-pathway targets identified for all scaffolds can be found in Fig. S2. Many off-pathway targets, including 1XQ8 and 2KKW above, are themselves deposited structures of monomeric *α*-synuclein. Fibril PDB entries of *α*-synuclein are excluded from the off-pathway target criterion calculation to avoid double counting the contribution of fibril to the ranking. Of the many *α*-synuclein fibril structures in the PDB, the only one identified as an off-pathway target by the above cutoff criteria was 2N0A, which happens to be the PDB entry we used to generate the fibril ensemble and perform epitope prediction. On the other hand, the structured monomer entries still contribute to the off-pathway target criterion because they are distinct from the simulated monomer ensemble. These monomer PDB entries are partly structured (containing *α*-helices) by binding to micelles (e.g. PDB 2KKW and 1XQ8), and are thought to be involved in various aspects of *α*-synuclein physiology (*37* –*42*). Thus, membrane-bound PDB monomer structures are treated separately from the unstructured isolated monomer ensemble.

### 2.6 Cyclic peptides with predicted conformational similarity to experimentally determined fibrils most effectively seed fibril aggregation

Conformational cyclic peptide scaffolds showed seeding activity in ThT aggregation assays (Fig. 7). Cyclic peptides that were predicted to have the highest conformational similarity to fibrils showed the most effective seeding activity. This supports the ranking by conformational similarity method proposed here. The two replicates of seeding experiments in Fig. 7 showed dramatically different patterns of the aggregation. The data for both runs were well-fit by an aggregation model including primary nucleation, elongation, and fragmentation (Fig. S4). The rapid upswing of the aggregate concentration in run2 is due to a much larger global fragmentation rate than run1 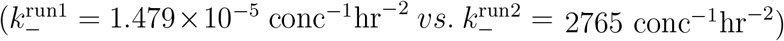.

**Figure 7:**
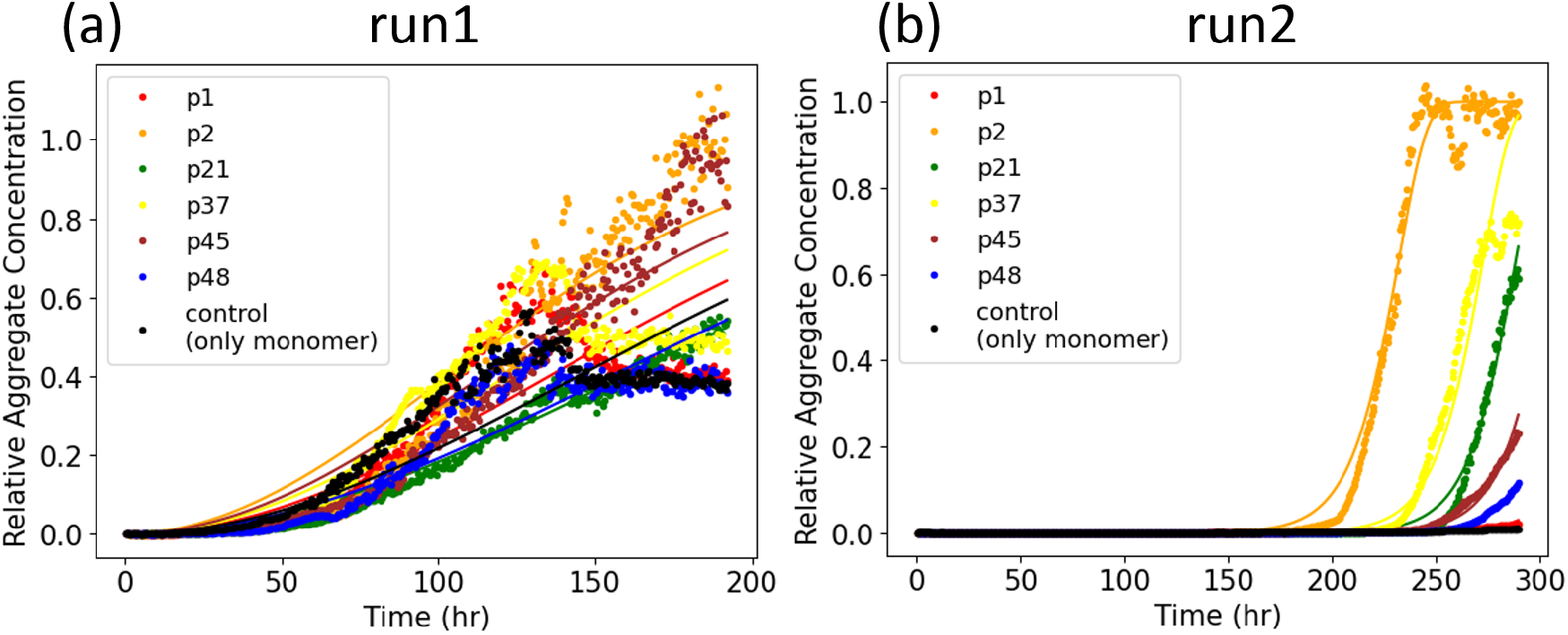
Two replicate experiments (run1 and run2) of seeded aggregation, as probed by ThT fluorescence. Curve fitting is obtained from the AmyloFit server (*43*).

Because the experimental assay is agnostic to *α*-synuclein fibril structure, the conformational similarity was investigated for multiple fibril structures of *α*-synuclein in the PDB. The embedding depth 𝒟_*f*|*c*_ of multiple PDB fibrils in each cyclic peptide ensemble is calculated. Different cyclic peptides may seed different fibril structure morphologies or strains.

We also assume the nucleation rate as determined by the model fitting may vary between different replicates (experimental runs), due to a number of experimental factors including evaporation and low signal for run 1, as mentioned in the Methods, as well as the possibility that a different fibril morphology may drive aggregation in each replicate.

We calculate the embedding depth 𝒟_*f*|*c*_ of 32 *α*-synuclein fibril structures deposited on the RCSB PDB and containing the sequence motif “EKTKEQ”. The procedure of the depth calculation is the same as treating each fibril PDB as an off-pathway target. Embedding depth is averaged over all instances of the epitope in each chain (typically 2 or 3 depending on the sequence), and over all chains in the fibril. Table S2 summarizes the values of 𝒟_*f*|*c*_ between each of the 32 fibril structures and each of the cyclic peptide ensembles for the 6 cyclic peptides examined in the seeding assay. The depth of the PDB fibrils in the monomer (the last column in Table S2) is also calculated, which provides the background nucleation propensity for all the cyclic peptide seeds.

We readily notice from Table S2 that the embedding depth of fibrils in peptides P21 and P48 is essentially zero, and these are also the worst nucleating seeds in the assay in run1, and are the worst and 2nd worst seeders on average. We also notice that the embedding depth of fibrils in peptide P2 is clearly the highest, and it is also the best seeder in the experiment. This conclusion is robust across both replicates of the experiment, despite variability in other aspects such as nucleation rates (Fig. S4).

When averaged over all fibrils, (𝒟_*f*|*c*_) shows good agreement with the nucleation rate, especially in run 1 (Table S2). This suggests the possibility that multiple fibril morphologies or strains may be seeded by the cyclic peptides, commensurate with their embedding depth. From the embedding depth analysis in Table S2, fibril structures that may be predicted to perform better at seeding aggregation can be identified as having a monotonic increase in 𝒟_*f*|*c*_ in the same rank order as the increase in observed nucleation rate *k*_*n*_. We may also predict a fibril structure is a reasonably good seeder if it has no more than 1 violation of this increasing trend. We found that run1 and run2 shared three of these candidate seeding structures (6OSM, 6CU7, and 6OSJ). This number does not reach statistical significance however (the expected number of shared structures under the null hypothesis is 2.18; pvalue of having 3 shared better seeder is 0.090).

As mentioned above, the two replicates of the experiment (run 1 and run 2) had different degrees of evaporation rate and signal, and showed variability in nucleation rates and global fragmentation rates. However, both had the same positive (P2) outlier (Fig. S4). In addition to experimental variability, this may also be due to different fibril morphologies that were nucleated stochastically in each of the experiments. As well, expecting our similarity metrics to perfectly explain the seeding experiment would implicitly assume that the cyclic peptides should act as on-pathway fibril seeds. However, this property is not equivalent to the ranking criteria that were used above. This discrepancy is discussed further in Section 2.10 below. The imperfect correlation between predicted fibril conformational overlap and observed nucleation rates in these experiments may be due to the fact that the cyclic peptides can also template off-pathway oligomer formation, which could frustrate the formation of fibrils, leading to lower fluorescence signal (*14*). Further experiments extending the preliminary data in this pilot study would be necessary to elucidate the molecular seeding mechanisms at play.

The criteria in the assay for seeding aggregation are distinct from the criteria for ranking cyclic peptides as immunogens for oligomer-selective antibodies. The above seeding experiment is thus not an ultimate proof of the ranking method described in this paper, which would be the successful production of conformation-specific antibodies using the computational methods to select top candidates from a pool of candidate antigen-presenting peptides. Rather, the analysis of the seeding experiment provides a plausible explanation of the seeding activity through the same similarity metrics used in the glycindel ranking.

### 2.7 Weighting the importance of ensemble properties

Ranking the peptide “glycindel” scaffolds here involved extensive conformational analysis and a ranking procedure with weighted criteria based on this analysis. Assigning appropriate weights for each criterion is a non-trivial task, which depends on biological intuition, and analyzing the consequences arising from screening criteria with a given set of weights. For this specific ranking, we put twice the weight on dissimilarity from the fibril and the monomer than similarity to the stressed fibril. We chose this weighting scheme because the stressed fibril ensemble generation process contains more prior assumptions, in which we hypothesize that a partially disordered fibril ensemble is enriched in oligomer-selective conformational epitopes.

The fibril ensemble is obtained by generating the “native basin” ensemble of the solidstate NMR resolved structure (PDB entry 2N0A). The monomer ensemble is obtained by generating disordered conformations of an isolated monomer of *α*-synuclein. The confidence in the accuracy of the fibril ensemble and monomer ensemble are greater than that of the stressed fibril oligomer model ensemble. The off-pathway target criteria have a slightly higher weight than the similarity of stressed fibril, because minimal off-pathway target effects are desired and are key feature of monoclonal antibody therapy.

### 2.8 Other possible scaffold design strategies

The *in silico* screening method introduced in this paper can generally be applied to various different scaffolding methods. There are many other scaffolding methods that have been previously developed (*44* –*48*) besides cyclizing the epitope with a variable number of glycine residues. Apart from these methods, several possible extensions of glycindel cyclic peptide scaffolding method may be directly implemented. For example, proline can be used for scaffolding in addition to glycine, since it is relatively inert chemically compared to other amino acids, and it is also able to constrain the cyclic peptide ensemble by adding conformational rigidity (*49*), which may be exploited to bias the scaffolded ensemble more effectively towards the stressed fibril ensemble. Large-scale computational design, which explores a vastly larger phase space of possible sequences, may also be implemented, e.g. through Rosetta (*44, 50*). We have not pursued these design methods here, mainly because of the criterion to avoid antigenic sequence outside of the epitope of interest.

While not common, epitopes of disease-specific antibodies may occasionally involve two or more requisite, discontiguous segments along the primary sequence of the peptide chain. Such a scenario arises, for example, in the anti-tau antibody zagotenemab, whose murine precursor was raised from purified paired helical filaments as an immunogen (*51*). Selective presentation of discontiguous epitopes can be facilitated by structural fixation of peptides, e.g. using cysteine–benzyl bromide chemical linkages (*52*). The antibodies raised by cyclic peptide glycindel scaffolds would not have this property. Many other disease specific anti-bodies in clinical development for Parkinson’s disease (*53*) as well as other neurodegenerative diseases (*1*) similarly target contiguous epitopes.

Cyclic and multicyclic peptide mimics of antibodies have been developed to help resolve the challenges of high production cost, structural stability issues, and low cellular uptake efficiency (*54* –*56*). Cyclic peptide loop length has also been varied and performance compared between 3 cyclic peptide constructs, to best mimic an anti-idiotypic/antireceptor antibody (*57*), and cyclic peptide mimics of transthyretin (TTR) have shown cyclization and loop length dependence in their efficacy to block A*β* aggregation (*15*). The conformational similarity methods and ranking method that we have developed here should have direct application to these areas of research. Cyclic peptide ensembles can be constructed *ab initio*, while collecting ensembles for larger protein complexes (e.g. an antibody with given paratope sequence) would typically require an experimentally resolved structure or reliable prediction such as an AlphaFold model (*58*).

Different carrier proteins that conjugate to the scaffolded epitope such as nanoparticles or lipid micelles have also been successfully implemented previously (*59* –*62*). Similarly, a linear peptide epitope of CGTKEQGGGG conjugated to a much larger carrier can be simulated as a peptide with cysteine SG atom spatially fixed at *x* = 0, and a boundary constraint such that it is forbidden to access the half-space region *x <* 0. Such a peptide in fact exhibits desirable ensemble overlap properties (𝒟_ensemble-in-monomer_=0.142, 𝒟_ensemble-in-stress_=0.207, 𝒟_ensemble-in-fibril_=0.000). Thus, conjugating on a non-interacting surface might be sufficient to modify the ensemble of the epitope to mimic the model oligomer.

The above possible extension strategies are interesting future tests for computational and experimental studies.

### 2.9 Previous applications of glycindel scaffolds

The glycindel scaffolding method has been applied to raise oligomer-selective antibodies to A*β* in a previous study (*23*), though we did not pursue a systematic ranking or optimal glycindel in this previous work. In Silverman et al. (*23*), an A*β* epitope containing the primary sequence SNK and predicted to be selectively exposed on oligomers (*63, 64*), was glycindel-scaffolded into two cyclic peptides: Backbone cyclized cyclo(CGSNKGG) and disulfide-bond cyclized cyclo(CGSNKGC). Ensemble similarity analysis of the two cyclic peptides showed low overlap with both A*β* fibril and monomer ensembles (*23*), and antibodies raised to the cyclic peptides exhibited oligomer-selectivity. Similarly, in another previous work (*24*), a predicted oligomer-selective epitope on A*β* (*63*) containing primary sequence HHQK was glycindel-scaffolded using cyclo(CGHHQKG), which when used as an immunogen could also raise an oligomer-selective antibody. A systematic test of the screening method described in this work has not yet been performed; It would involve immunization and antibody generation from both low-ranked and high-ranked glycindels, and a successful test of conformational selectivity wherein oligomer-selective antibodies arose from highly-ranked glycindels but not low-ranked glycindels.

In currently ongoing work, antibodies have been raised to *α*-synuclein using four highly-ranked glycindel peptides, cyclo(CGGGEKTKGG), cyclo(CGGGGEKTKGG), cyclo(CGTKEQGGGG) and cyclo(CGGTKEQGGGG). These antibodies show conformational selectivity toward *α*-synuclein oligomer and soluble pre-formed fibrils, while sparing healthy monomer. The above proposed glycindels are not the top-ranked ones in this study however, since at the time the antigens were selected, the embedding depth as a screening metric was not developed. The success of these glycindels based on more preliminary screening criteria implies some plasticity in the ensembles explored by the glycindels, and thus likely some leniency in selecting the best candidates.

### 2.10 Evaluating the assumptions of the collective coordinate prediction of the EKTKEQ epitope in *α*-synuclein

There are several alternative *α*-synuclein fibril structures with different polymorphs deposited in RCSB protein data bank (PDB) (www.rcsb.org) that we could have used for this study (e.g. PDB entries 6CU7, 6CU8, 6H6B and 6FLT), which raises the question of whether the same epitopes would have been predicted with these alternative structures. There is a precedent for some oligomer-selective epitope commonality among polymorphic A*β* fibril structures (*63*). The single protofilament solid-state NMR structure of the *α*-synuclein fibril (PDB ID: 2N0A) used here for epitope prediction is similar to both the dominant rod (PDB ID: 6CU7) and twister (PDB ID: 6CU8) polymorphs for 38 matched residues of the *α*-synuclein fibril structures, with an RMSD of 3.5Å and 3.8Å, respectively (*65*). Another cryo-EM structure of a truncated *α*-synuclein fibril, which includes the first 121 residues (*66*), has also been shown to have structural similarity to the rod polymorph with an RMSD of 2.1Å (*65*). These similarities between the aggregating units of different polymorphs of *α*synuclein suggests that there may be some commonalities in collective coordinate-predicted epitopes. That said, the fact that these other fibril structures did not appear as strong hits in our off-pathway analysis suggests that if there was overlap in the exposed epitopes, the oligomer conformational ensemble would not be well-modelled by any of the cyclic peptides we investigated. Systematic analysis of the predictions for these structures would have to be done to validate this hypothesis, and is a topic for future work.

In the context of Alzheimer’s disease, both on-pathway or off-pathway oligomers have been associated with A*β*-derived toxicity (*67* –*69*). A similar situation arises in Parkinson’s disease, wherein both oligomeric and fibrillar species have been associated with cytotoxicity (*70*). Some studies have shown that at least some off-pathway oligomeric species are relatively inert, and that oligomers conducive to fibril formation are cytotoxic (*71*), while other studies have shown that small molecules can alter the conformations of off-pathway *α*-synuclein oligomers to species that are non-toxic (*72, 73*). The issue of optimizing therapeutic strategies by targeting on-pathway or off-pathway oligomers is an important one that is an area of current active research.

Using a stressed fibril for the prediction of oligomer-selective epitopes may appear to imply that the predicted epitopes are present in conformations that are on-pathway to fibril formation. However, the main assumption in the collective-coordinates prediction method is only that those segments of primary sequence that are prone to exposure in a stressed fibril will also be prone to exposure (and thus antibody accessible) in an oligomer. To investigate the generality of such collective coordinate-predicted epitopes, we examine the solvent accessible surface area (SASA) as a function of sequence for 5 *α*-synuclein fibril polymorphs chosen based on their structural dissimilarity (Fig. 8g). The SASA as a function of sequence index is plotted in Figs. 8a-e, and the region consisting of sequence index 57-62, corresponding to the collective-coordinates-predicted epitope EKTKEQ, is highlighted in the figures. The SASA averaged over all 5 structures is also plotted in Fig. 8f, where it can be observed that the 6 amino acid epitope region has the largest SASA of all such regions across the primary sequence.

**Figure 8:**
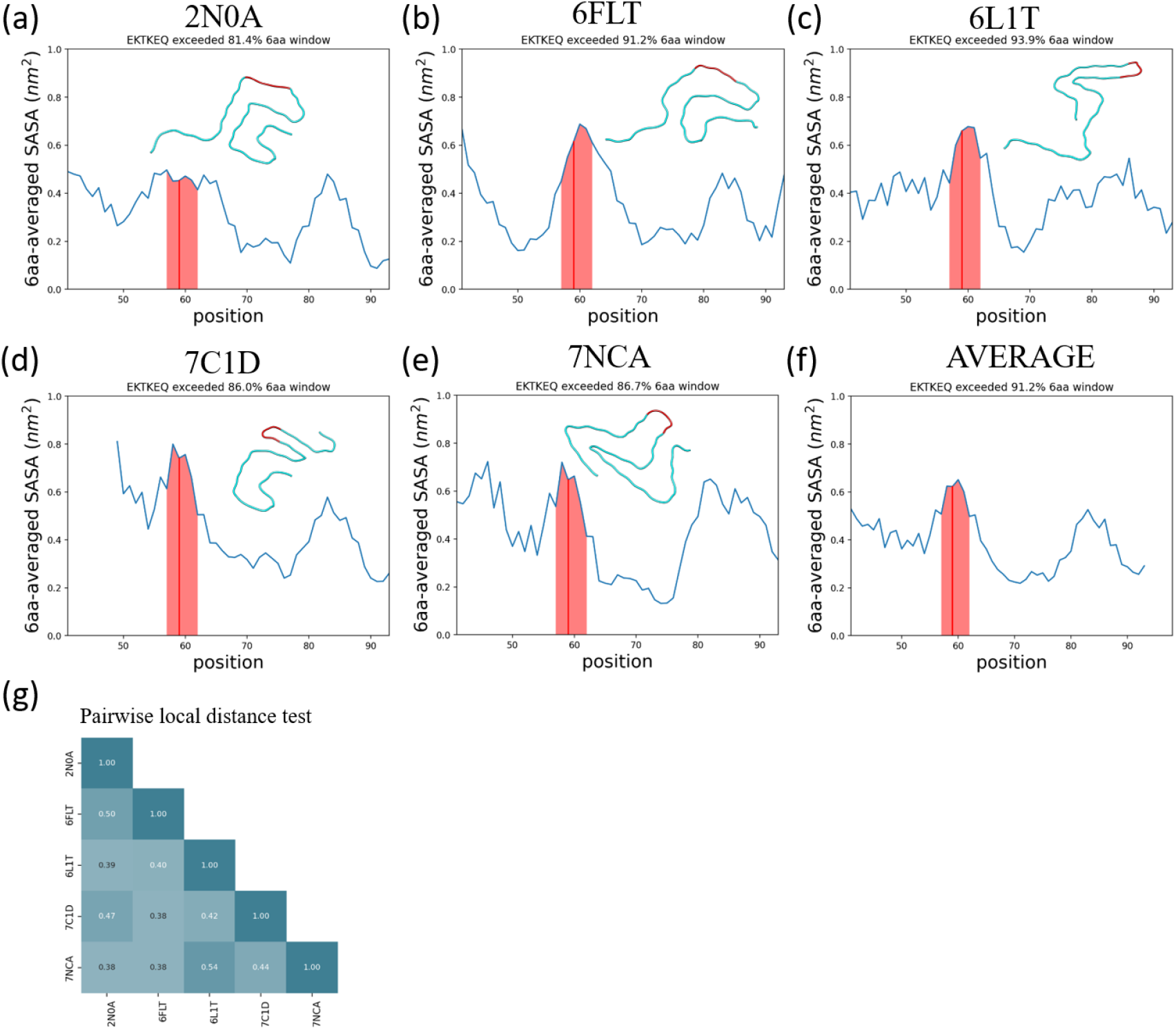
The *α*-synuclein epitope, EKTKEQ, is found to be highly exposed in five structurally distinct fibrils. (a-e) Averaged SASA as a function of residue position. A rolling average window of 6 amino acids was applied. The window that contains EKTKEQ (residues 57-62), as indicated by the red line, exceeds more than 80% of the other windows in all fibril structures analyzed. The shaded region contains the rolling average values for residues 57-62. In each panel, a single chain of each fibril structure is aligned and rendered to show their structure heterogeneity. (f) The average SASA across all 5 fibrils. The epitope region has the highest average SASA across the whole structured sequence. (g) The pairwise local distance test (lddt) (*74*) shows that the analyzed fibrils are all mutually dissimilar.

**Figure 9:**
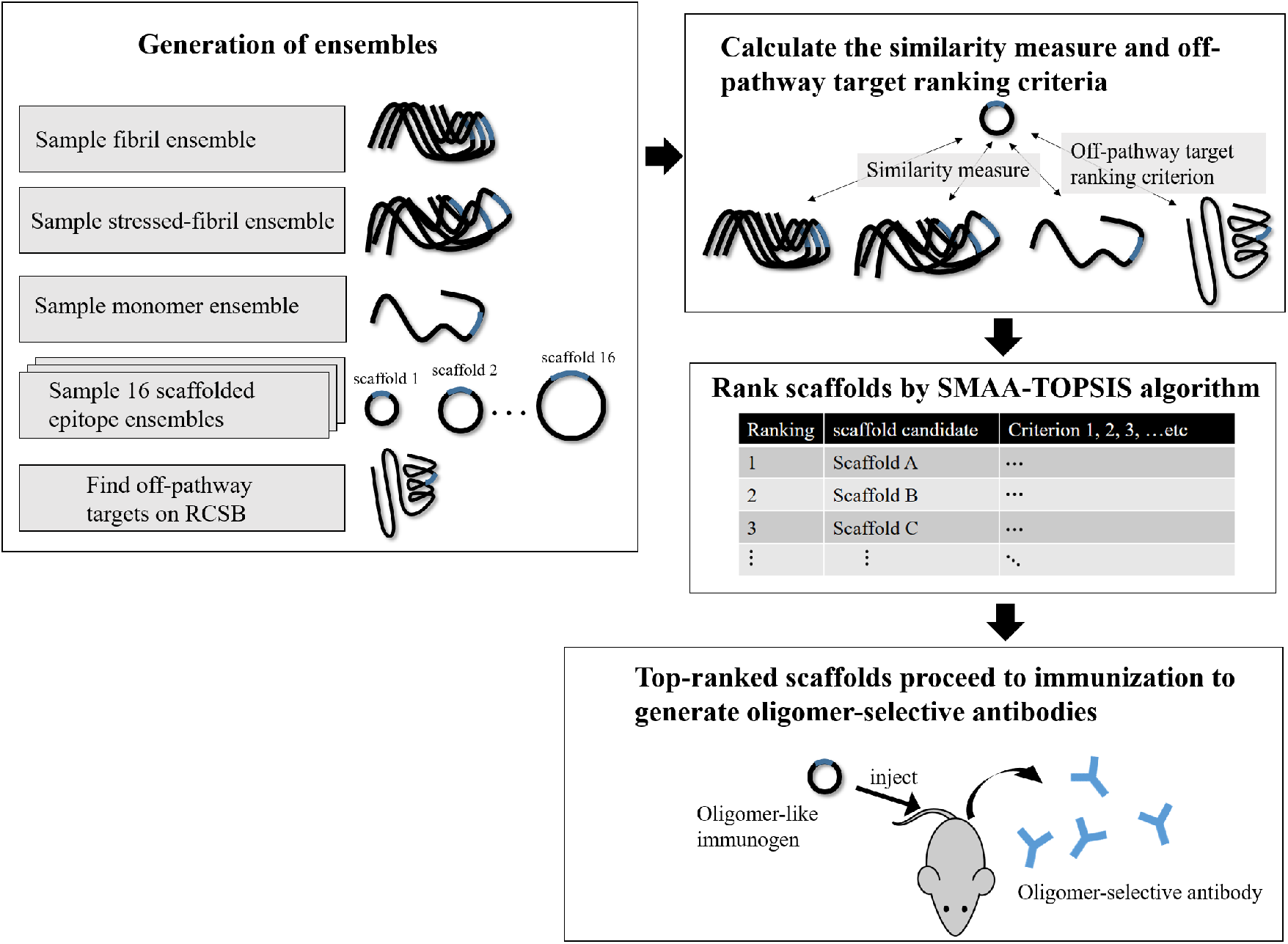
The workflow of *in silico* screening.

**Figure 10:**
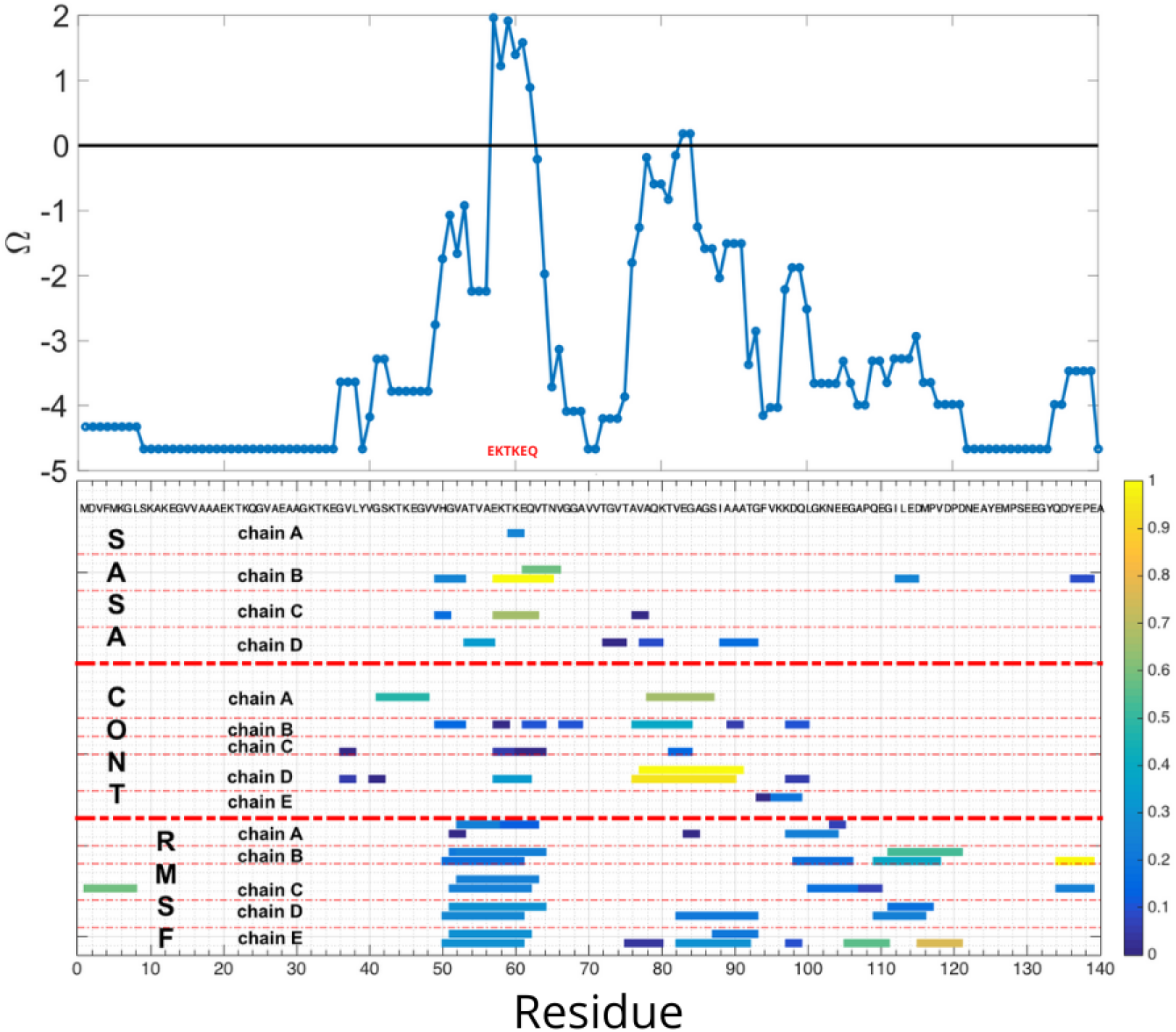
Collective coordinate epitope prediction for *α*-synuclein, using three criteria of increased SASA, loss of native contacts, and increased fluctuations (RMSF). Several epitopes were predicted by each criterion; however, only a single consensus epitope EKTKEQ was predicted. Chain E is not shown for ΔSASA because no epitope is predicted.

Because distinct fibril morphologies are not observed to readily convert due to large interconversion barriers (*70*), the above result implies that the epitope is either a.) A generic on-pathway to fibril epitope, which happens to be largely independent of the particular aggregated structure, or b.) A generically exposed region of aggregated structures, oligomer or fibril, based on physico-chemical grounds of charge, hydrophilicity, and weakly stabilizing interactions. The collective coordinates-predicted epitopes may be more broadly applicable than the fibril structure from which they were derived, and antibodies to them do not necessarily select for conformations on-pathway to the fibril. (*23, 24*) In the context of natively folded proteins, antibodies to collective coordinates-predicted epitopes of SOD1, which misfolds and aggregates in SOD1-related ALS, do not target native protein or nearly folded variants, but rather are selective to pathological inclusions.

### 2.11 The role of glycindels in therapeutic development pipelines

The *in silico* immunogen screening method developed here provides an additional route to aid the development and optimization of a peptide immunogen that can be faster and cheaper than experiments. Experimental screening methods can then be performed on a subset of top-ranked scaffolds to save available resources. Some examples of downstream *in vitro* screening experiments are Thioflavin T (ThT) aggregation assays (*24, 75, 76*) (see

Section 2.5), surface plasmon resonance (SPR) assays to measure binding affinity to antibodies and conformational selectivity (*27*), and Förster resonance energy transfer (FRET) assays (*77*) to measure aggregation tendency, since scaffolds that can trigger aggregation of normal *α*-synuclein may have more contribution to the pathology because of greater seeding propensity. These *in vitro* assays are typically followed by various *in vivo* studies as part of pre-clinical development. The ability to use computationally generated conformational ensembles as a screening method for candidate immunogens can aid and accelerate therapeutic development.

## 3 Methods

### 3.1 *α*-synuclein epitope prediction

In our approach, we operate from the hypothesis that a partially disordered protofibril ensemble is enriched in oligomer-selective conformational epitopes. That is, similar regions are exposed in both toxic oligomers and stressed protofibrils. If so, the stressed fibril may be used to predict oligomer-selective epitopes. This hypothesis is supported by previous *in vitro* and *in vivo* evidence (*23* –*25*).

The EKTKEQ epitope in *α*-synuclein was predicted by the Collective Coordinate (CC) algorithm, which has been used previously to predict misfolding-specific epitopes in superoxide dismutase 1 (SOD1) (*25*). The CC algorithm predicts epitopes by identifying local unfolding events when a global denaturing stress is applied, where regions of a protein or multi-protein aggregate deviate structurally from their native structural conformation. Three metrics are used to measure local disorder: increased solvent accessible surface area (ΔSASA), loss of native contacts (ΔQ), and increased root mean squared fluctuations (ΔRMSF).

The procedure for predicting an epitope involves implementing a global unfolding potential, which biases the system to have 0.65 of its total native contacts. Ten independent biased ensembles of an *α*-synuclein fibril structure (PDB ID: 2N0A) using the CC algorithm, as well as a single “native basin” ensemble of the fibril using molecular dynamics (MD) simulation (*25*), are generated to calculate ΔSASA, ΔQ and ΔRMSF. The “native basin”ensemble serves as a reference for calculating the above three difference values. Multiple independent biasing simulations were performed in order to ensure consistency in the regions of the fibril structure that are observed to have relatively higher values of ΔSASA, ΔQ and ΔRMSF, thereby avoiding predictions based on rare fluctuations that might be present in a single biased simulation.

Fig. 10 bottom panel shows the sequence motifs larger than 3 amino acids that are predicted as epitopes by each of the three metrics, for each of the 5 chains in the protofibril structure (PDB 2N0A). Several epitopes were predicted by each metric; however, the consesus epitope EKTKEQ is predicted by all the three metrics, and was taken as the final predicted epitope for *α*-synuclein (Fig. 10 top panel).

### 3.2 Scaffolding of epitopes

We scaffold epitopes by constructing cyclic peptides containing both the epitopes and a variable number of glycines. We refer to the resulting scaffolds as glycindels. First, the predicted epitope (EKTKEQ), or a shorter sequence that is subsumed by the epitope such as EKTK, is flanked on both sides with consecutive glycines. This is implemented computationally by mutating the native flanking residues to glycine using SCWRL4 (*78*). In addition, one more residue is mutated to cysteine on the N-terminal side of the sequence. Although, in principle, other amino acids or a combination thereof can be utilized, our choice of using glycines helps to focus immunogenic effects to just the epitope, since glycine is relatively chemically inert compared to other amino acids (*79*), and has minimal immunogenicity (*80*). The cysteine is present to conjugate the peptides to immunogens such as Bovine serum albumin (BSA) or Keyhole limpet hemocyanin (KLH) via a disulfide bond, increasing the likelihood of generating antibodies to the scaffolded epitope. The topology of the cyclic peptide is obtained by head-to-tail linkage of the termini using a locally written Python script (https://github.com/PlotkinLab/Backbone-linkage-of-cyclic-peptides).We refer to a cyclic peptide epitope with *n* N-terminal glycines and *m* C-terminal glycines as cyclo(*C*-*G*_*n*_-(Epitope)-*G*_*m*_) or (*n, m*)Epitope. The cyclized peptide is then energy minimized in GROMACS using steepest descent algorithm, before running equilibrium MD simulations.

### 3.3 Generation of ensembles

Generally, we collect four different equilibrium molecular dynamics (MD) ensembles for each epitope. These correspond to the epitope in the context of an isolated monomer, the fibril, the stressed fibril oligomer model, and the cyclic peptide scaffold. We perform MD simulations with the open-source GROMACS (*81*) package and the community developed PLUMED library (*82*). The force field used here for all ensemble sampling is CHARMM36m (*83*), which is a modified version of CHARMM36 with improved modeling for disordered proteins. Each epitope ensemble is obtained specifically as follows.

#### 3.3.1 Fibril ensemble

The conformational sampling starts from an existing experimentally resolved *α*-synuclein structure determined by solid-state NMR (*84*) (PDB ID: 2N0A). A protofibril composed of 5 chains (chains A-E of 2N0A) is solvated in 150mM Na and Cl aqueous solution, such that the system is neutral. After 50ps constant volume (NVT) and 150ps constant pressure (NPT) equilibrium simulations under positional restraints on heavy atoms, we perform an equilibration MD simulation up to 100ns until the convergence is seen from RMSD during the simulation. A 20ns equilibration MD continued from the previous 100ns simulation is then performed, for collecting the “native basin”ensemble of the fibril, from which we sample snapshots at 20ps intervals. Only configurations of the middle chains (chain B-D) are collected in the ensemble, in order to reduce edge effects in the simulation. In total, the regularly spaced sampling results in 3003 configurations in the fibril ensemble obtained by this procedure. The above procedures of solvation and NVT-equilibration and NPT-equilibration are also implemented for the other ensembles described below, before the MD production runs.

#### 3.3.2 Stressed fibril ensemble

The stressed fibril ensemble is a partially disordered fibril ensemble used to predict conformational epitopes similar to what might be presented by the oligomer. To generate this ensemble, a time-dependent global bias potential *V* (*Q, t*) is implemented to partially unfold the *α*-synuclein fibril, where

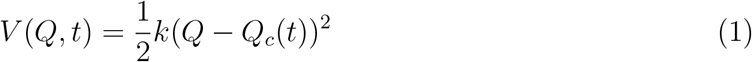

Q in equation (1) is a collective coordinate defined as the normalized count of the native contacts (*25*): Native contacts are pairs of heavy (non-hydrogen) atoms within 4.8Å of each other that are in different amino acids labeled by primary sequence residue index *α, β*, that satisfy |*α − β*| *≥* 3, and persist over 5 percent of the time in the first 100ns of equilibrium fibril simulation. Formally, the collective coordinate Q for any structure characterized by a set of heavy atom distances, *r*_*ij*_, is defined as follows:

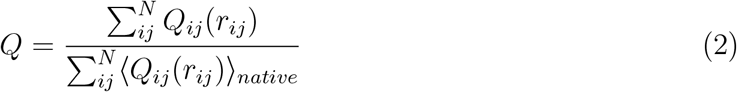

where

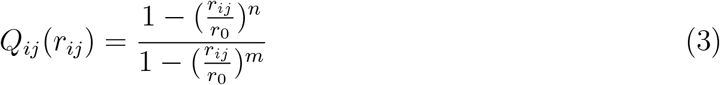

where we take *r*_0_ = 4.8Å, *n* = 6 and *m* = 12. Equation (3) approaches a step function for large *m* and *n* (for *m > n*), but is smooth and differentiable (*25*), which is a desirable property for the contact function. A contact function *Q*_*ij*_(*r*_*ij*_) is defined for each heavy-atom pair *i, j* in the list of native contacts, which rapidly goes to one when *r*_*ij*_ is slightly lower than *r*_0_ and rapidly goes to zero when *r*_*ij*_ is slightly larger than *r*_0_. The quantity in the numerator of Q in equation (2) is the sum of *Q*_*ij*_ in an arbitrary structure, where 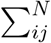 is summation over the native contact list, and the quantity in the denominator is the Boltzmann average of the *Q*_*ij*_ in the fibril “native basin” ensemble. Thus, Q is typically a number between zero and unity.

*Q*_*c*_(*t*) in equation (1) stands for the target value of the collective coordinate, Q. It is a time-dependent quantity that starts from a value corresponding to the counts of all native basin contacts in the fibril metastable equilibrium ensemble (*Q*_*c*_ = 1), which is then taken to linearly decrease with time, from *Q*_*c*_ = 1 to *Q*_*c*_ = 0.65 over 50 ns. Afterward, the bias is held fixed at *Q*_*c*_ = 0.65 for 150ns. During the last 50ns of this second period when *Q*_*c*_ is held fixed at 0.65, snapshots from the stressed fibril ensemble are collected at an interval of 15ps. The above process is repeated 10 times, including the ramp down and subsequent biased equilibration and sampling, to average the stochastic unfolding process of the protofibril and obtain a reliable biased ensemble. The capping chains at the ends of the fibril are discarded to avoid edge effects, yielding a total ensemble consisting of 50*ns/*15*ps ∗* 3 chains *∗* 10 repeats = 100000 sampled structures. A subset with 3500 configurations is randomly sampled from the total structural ensemble for efficiency of the calculation. In order to obtain an ensemble of the epitope that is partially disordered and exposed to solvent, we add on additional constraint on SASA of the epitope in the ensemble, as follows. Among the 3500 configurations, we further discard those structures that have a lower epitope SASA than the equilibrium epitope SASA, generating a stressed fibril ensemble with exposed epitope consisting of 3407 configurations. We note that at this point the epitope has already been predicted, and the above calculations are for comparison of the stressed/exposed protofibril ensemble with cyclic peptide scaffold motifs.

#### 3.3.3 Monomer ensemble

Since *α*-synuclein is an intrinsically disordered protein (IDP) (*85*), its equilibrium ensemble is relatively difficult to sample sufficiently by normal MD simulation (*86*). We have previously developed a method for generating equilibrium ensembles for IDPs (*87*), and have applied this method to generate equilibrium ensembles for several IDPs such as *α*-synuclein, A*β* peptide, and prothymosin *α*. We have also used the method to model the unfolded ensembles of natively folded proteins, including disulfide bonded unfolded ensembles such as that for SOD1 (*87*).

Here, we modify the method in reference (*87*) (which included a coarse-graining step) to generate an equilibrium monomer ensemble for *α*-synuclein. We do not employ a coarsegraining step. The steps involved in our ensemble generation include: **1.)** Generation of a conformationally diverse ensemble via the pivot algorithm and a “crankshaft” move wherein two randomly selected end points are fixed (*87, 88*), and **2.)** Equilibration of each pivot– and crankshaft–randomized structure for a short simulation time. The descriptions for these steps follows.

The initial *α*-synuclein monomer configuration is obtained by extracting a single chain from the fibril structure (PDB ID: 2N0A). This structure is then altered by employing a generalization of the pivot algorithm (*89*–*92*), which is an efficient algorithm for generating ensembles for a self-avoiding random walk, as well as a “crankshaft” move (*87, 88*) that randomizes *φ* and *ψ* angles between two randomly-selected backbone atoms along the peptide chain, such that the randomly-selected backbone atoms remain fixed. Pivot moves and crankshaft moves are attempted with equal probability. If a move results in a steric clash, it is rejected, and the next randomly selected move is then attempted. We take a simple approximation wherein *φ* and *ψ* are sampled from a (2*π* periodic) von Mises probability distribution with mean 0 and variance 1, i.e. e^cos *φ*^*/*(2*πI*_0_(1)), where *I*_0_ is the modified Bessel function of order 0. Pivot/crankshaft moves are repeatedly attempted until there are *N* successful pivot/crankshaft moves, where *N* is the length of the chain. This process completely randomizes the conformation such that information about the original structure is lost.

After a randomized conformation is obtained, the structure is then solvated in a box determined by the principal axes wherein the peptide is at closest 1.2 nm from the box edges. The structure is then energy minimized, equilibrated in explicit modified TIP3P water and 150mM salt in an NVT ensemble for 100ps, and then equilibrated in an NPT ensemble for 300ps. This process generated 1587 different structures as initial configurations. We then performed a 3ns equilibrium simulation starting from each of the above initial configurations, collecting a snapshot every 1ns to be added to the monomer ensemble. As a result, we obtained a monomer ensemble with 4643 configurations (i.e. 93 simulations did not reach 3ns and 16 simulations did not reach 2ns within the simulation wall-clock time, due to variable simulation box sizes). IDP ensemble generation is an active area of research and several other methods of computational ensemble generation have been developed (*93*).

#### 3.3.4 Scaffolded epitope ensembles

The construction of the initial structure used for scaffold simulations is described in Methods Section 3.2. After minimization, solvation, NVT-equilibrium and NPT-equilibrium simulation, a 300ns MD simulation is carried out, from which an initial ensemble is collected with constant sampling interval of 40ps. Since the cysteine side chain has to be solvent-accessible to form a disulfide bond with the carrier protein (KLH or BSA), some configurations were discarded if the SG atom in the cysteine was buried or if the sidechain of the cysteine pointed inside the cyclic peptide: The SG atom is defined as buried if its SASA is lower than 70% of that in an isolated Gly-Cys-Gly tripeptide (80Å^2^). The cysteine side chain is defined as inward pointing if its dihedral angle *ψ* (angle between CB atom and the backbone plane) is within [-90, 90] degrees. In practice, the scaffolded cyclic peptide ensembles thus processed have a different number of configurations for each cyclic construct, ranging from 2352 to 5468 configurations. The fraction of the ensemble that satisfied the above criteria varied from about 39% to 97%.

### 3.4 Projecting ensemble distributions to lower dimension

We first let *N* be the sum of the number of structures in all four ensembles: Monomer, fibril, stressed fibril, and cyclic peptide scaffolds (Here *N* ranges from 13369 to 16485). We then construct a pairwise root mean squared deviation (RMSD) matrix that consists of the RMSD between any two epitope (EKTK, KTKE, or TKEQ) structures in any of the four ensembles. Each row of the *N × N* matrix contains the distances (RMSD values) from one structure. Two rows then constitute a 2-dimensional space giving the distances to two structures determined by which rows have been chosen. *N* rows, corresponding to the whole matrix, thus constitute an *N* -dimensional space where the location of each of *N* points, corresponding to each structure, is determined by the set of distances (RMSD) to all other structures (including itself). The set of *N* structures is thus represented by a distribution of points in an *N* -dimensional space.

Multidimensional Scaling (MDS) (*32*) or Stochastic Proximity Embedding (SPE) (*30*) was performed on the RMSD matrix to reduce the dimension from *N ×N* to *N ×D* where 𝒟 is the desired lower dimension. The value of 𝒟 depends on the application as described below.

As well, whether MDS or SPE is used depends on the application, and will be described in detail in Sections 3.5, 3.6, and 3.7 below.

Applying MDS or SPE results in an ensemble of points in a reduced number of 𝒟 dimensions. We then fit the effective distribution of the *N* points in 𝒟 dimensions using kernel density estimation (KDE). (*30*) Since KDE performance worsens exponentially with higher dimensional data sets as a result of the so-called “curse of dimensionality” (*94*), a dimensional reduction step is essential. A particular ensemble, e.g. the monomer structural ensemble, has a structural distance distribution in the RMSD matrix. This structural distance distribution has a corresponding KDE distribution of points. We refer to this KDE structural distance distribution as simply the structural distribution below.

### 3.5 Calculating ensemble overlap in 1D

We visualize the similarity between two ensembles intuitively by representing them in 1D (this is not used as a ranking criterion). The overlap is the percentage of a KDE distribution that overlaps with another KDE distribution. The formula we use here for the overlap is: Overlap 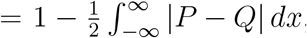, where *x* is the first MDS coordinate, and *P* (*x*) and *Q*(*x*) are two KDE distributions in this 1D coordinate. For example, for two gaussians of unit variance and mean separation *a*, the overlap is 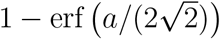.

### 3.6 Ensemble similarity measures

We compare conformational ensembles of the epitope in the four different contexts, including monomer, fibril, stressed fibril and cyclic scaffold, using two measures: Embedding Depth and Jensen-Shannon Divergence. These are defined as follows.

#### 3.6.1 Embedding depth

The embedding depth is most easily understood when taken as a measure between a single structure and a given ensemble. In this context, the embedding depth quantifies how deeply that structure is embedded within the ensemble, when both structure and ensemble are projected onto some coordinate metric such as MDS coordinates. The structure-to-ensemble embedding depth 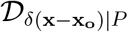, for a structure characterized by a point at **x** = **x**_**o**_ embedded in a distribution *P* (**x**), is defined as the fraction of the ensemble that has a lower KDE probability than the particular structure (Fig. 11):

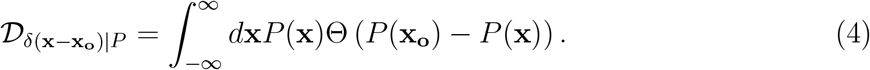

In equation (4), Θ(*P*) is the Heaviside step function, which returns 1 if *P >* 0, otherwise 0.

The mode of a distribution has an embedding depth of one because all of the ensemble has a lower KDE probability than the mode. On the other hand, outliers of a distribution have embedding depths close to zero.

**Figure 11:**
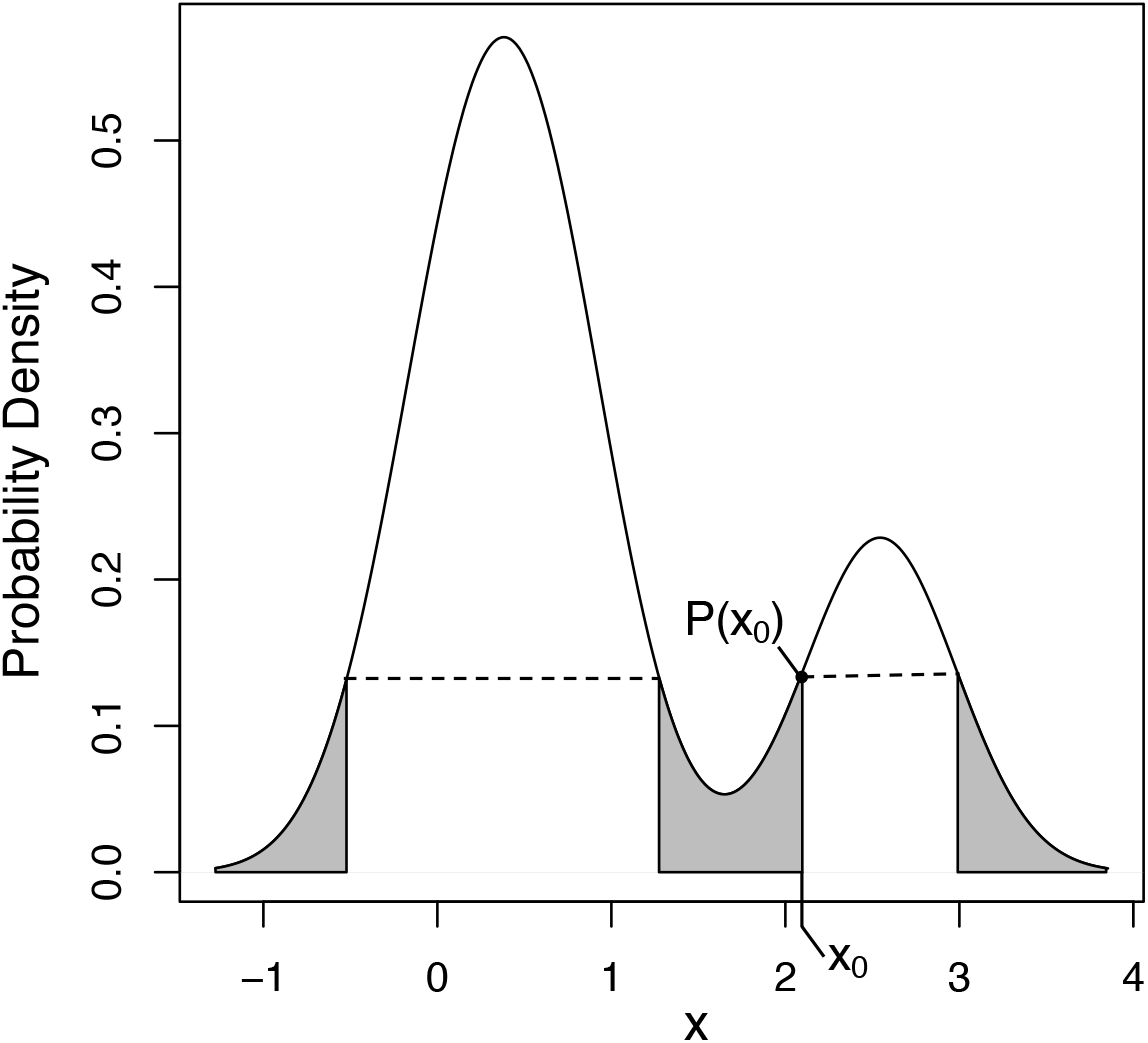
Illustration of the embedding depth of the point *x*_*o*_ in a multimodal distribution *P* (*x*). The embedding depth of point *x*_*o*_ is given by the integral over all parts of the distribution with probability less than *P* (*x*_*o*_). Note there are 4 points with the same *P* (*x*_*i*_) = *P* (*x*_*o*_) and thus the same embedding depth.

The embedding depth of one distribution *Q*(**x**) within another *P* (**x**) can be found by integrating equation (4) over the distribution *Q*(**x**):

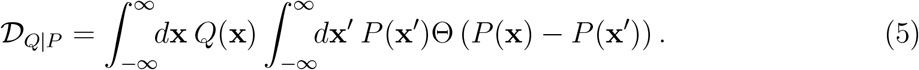

The embedding depth between two distributions is in general non-reciprocal, in that 𝒟_*Q*|*P*_ =*/ D*_*P* |*Q*_. It is straightforward to show that the embedding depth of a distribution within itself 𝒟_*P* |*P*_ = 1*/*2. This follows because the special case of 𝒟_*P* |*P*_ must be symmetric (reciprocal), so that switching **x** and **x** in 𝒟_*P* |*P*_ must yield the same equation. Therefore,

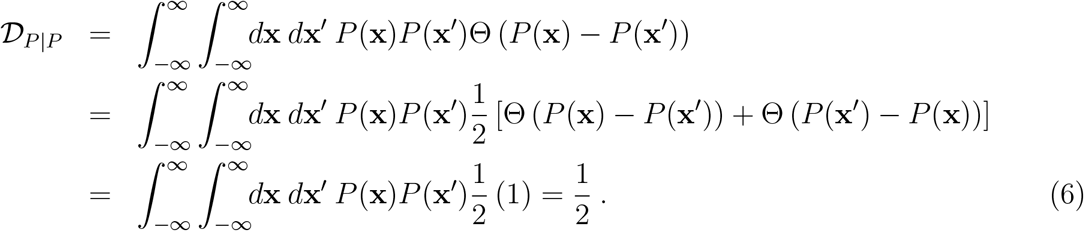

When used to compare two conformational ensembles, 𝒟_*A*|*B*_ or 𝒟_A-in-B_ represents the extent to which the ensemble *A* is subsumed by ensemble *B*. A similar concept in measuring the embedding of one ensemble in the other has been introduced before in the context of statistics (*31*), but we are not aware of such a measure being previously applied to conformational ensemble comparison. The embedding depth measures used for ranking (𝒟_cyclic-in-stress_, 𝒟_cyclic-in-monomer_, and 𝒟_cyclic-in-fibril_) are calculated in an MDS-reduced 3D conformational space, where 95% of the matrix information can be preserved (Fig. S1b).

Discrete, small sample size effects may also be considered for the embedding depth, however these effects are unlikely to arise in practice for conformational ensembles, which are typically large enough to achieve convergence.

#### 3.6.2 Jensen-Shannon Divergence (JSD)

The similarity between two different ensemble distributions is measured using the JensenShannon Divergence (JSD) (*29, 95, 96*), which is implemented here using the ENCORE software (*30*). JSD is a symmetrized and smoothed version of the Kullback–Leibler divergence (*97, 98*), 𝒟_*KL*_, which is a difference measure between two distributions. JSD is defined by

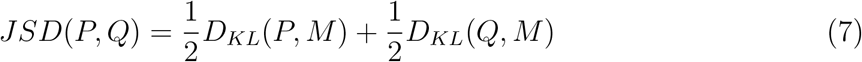

where

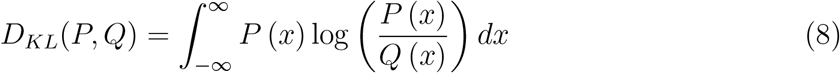

where *P* (*x*) and *Q*(*x*) are two conformational ensembles in SPE-reduced dimensional space, and *M* is defined by the average of the two distributions *P* and *Q*: *M* (*x*) = (*P* (*x*) + *Q*(*x*))*/*2. The value of the JSD measure depends on the dimensionality of the SPE-reduced space, with lower dimension tending to have smaller JSD (i.e. more overlap). The JSD values used for ranking cyclic peptide scaffolds (JSD_cyclic-fibril_, JSD_cyclic-monomer_, and JSD_cyclic-stress_) are produced by weight averaging the JSD from 3D to 11D by the inverse of the SPE residuals in each dimension (Fig. S1 for these residuals). By this measure, the higher dimensions contribute more to the weighted average JSD. The convergence test in Section S.2 shows that the information loss is about 3% in 3D and the information loss is less than 1% in 11D for scaffold (1,4)TKEQ for example (Fig. S1). JSD as calculated in equation (7) ranges from 0 to log(2), but in this paper, we normalize JSD to lie between 0 to 1.

### 3.7 Evaluating off-pathway target criteria

We search through the RCSB database of resolved protein structures, to find potential off-pathway targets of antibodies that could be generated by each cyclic peptide scaffold. An off-pathway target of concern meets three conditions: **1.)** same primary sequence as the epitope in the PDB structure, **2.)** structural similarity to scaffolded epitope, and **3.)** sufficiently high solvent exposure so that it is likely to be antibody-accessible.

The first criterion is assessed by finding “hits” when searching through the RCSB database (e.g. any protein that contains one or more TKEQ motifs). The second criterion is assessed by the PDB entry’s structural embedding depth in the scaffolded cyclic peptide ensemble. The embedding depth is calculated in the MDS-reduced 5D space of the RMSD matrix. The third criterion is assessed by the finding the fraction of the glycindel–scaffolded epitope ensemble with less solvent-accessible surface area (SASA) than the PDB entry (i.e. *f* (SASA_off-target-exceed-cyclic_)).

The severity of a potential off-pathway target for a given scaffold is determined by summing, over all PDB structure “hits”, the product of the structural embedding depth (𝒟_off-target-in-cyclic_) and the fraction of cyclic peptide ensemble SASA exceeded (*f* (SASA_off-target-exceed-cyclic_)). The criterion used for ranking (before a linear rescaling) is the negative of this severity, which is represented by equation (9) below:

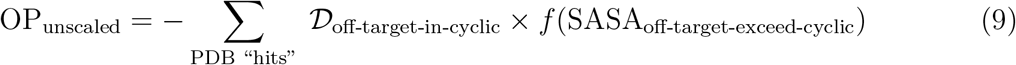

By this convention, a higher (less negative) criterion value corresponds to a more favorable scaffold. Since 𝒟_off-target-in-cyclic_ and *f* (SASA_off-target-exceed-cyclic_) are very small for most RCSB entries (i.e. most entries are not of significant concern), we employ a cut-off criterion wherein 𝒟 and *f* have to both exceed 5% for the entry to be included in a criteria calculation. Finally, the criteria are linearly rescaled to be within the range [0,1].

Sometimes, the epitope appears multiple times in a single RCSB PDB entry because of multiple deposited models, multiple chains or repetitive epitope occurrence on a single chain.

If such multiple occurrence happens, the 𝒟_off-target-in-cyclic_ and *f* (SASA_off-target-exceed-cyclic_) in equation (9) are taken as the average of the values over all epitope occurrences in the corresponding PDB entry.

The search result in the RCSB database will also contain the fibrillar and structured monomer *α*-synuclein entries. The fibril entries are excluded from the off-pathway target calculation, because we have already compared cyclic and fibril ensembles using JSD_cyclic-fibril_. We have also compared cyclic and unstructured monomer ensembles using JSD_cyclic-monomer_, however monomer structures in the RCSB database are kept because they are assumed to be at least partially structured due to effects such as peptide-membrane interactions.

Weaker binding of antibodies has occasionally been observed in epitope mapping by alanine scans, particularly in longer epitopes. This binding to single point mutants occurs more commonly in the peripheral regions of the epitope, and may generally be an important consideration. Our epitope lengths are only 4aa however, so mutation of one amino acid would be a significant 25% change in identity. We thus have not explored single point mutants in the off-pathway analysis here.

### 3.8 SMAA-TOPSIS ranking algorithm

Scaffolds are ranked using an adaptation of the so-called SMAA-TOPSIS algorithms, which stand for a combination of Stochastic Multi-criteria Acceptability Analysis and Technique for Order Performance by Similarity to Ideal Solution. (*99*) The ranking algorithm we employ here uses a cumulative measure of the retention probability that a candidate is retained in the top *n* leads (*27*). The method generates a ranking of candidates when multiple criteria are used for screening the candidates, and when those screening criteria have a unknown predictive importance *a priori*. The screening criteria as well as the importance weights for those criteria are both subject to error, and an exact weight assignment for the importance of the screening criteria is not needed. Instead, weight distributions are assigned, where the mean weight *w* represents the relative importance of each criterion and the standard deviation of the weight distribution represents the uncertainty in the importance for that criterion. The weight distribution is taken to be a uniform distribution:

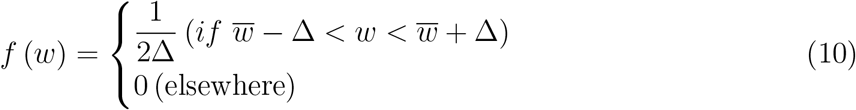

where *w* stands for the mean (or center) and Δ stands for the distribution width. All the criteria have Δ = 0.5 in this paper (or equivalently standard deviation *σ* of 0.288). The mean weights for each criterion are given in Table 1:

The ranking algorithm rescales all screening criteria to unity so weights can be properly applied. The Ideal Best and Ideal Worst entries are given similarity screening criteria of either all 1’s or all 0’s respectively; These numbers are given for each scaffold in Table S1. All 48 scaffolds show strong dissimilarity to fibril, whereas other screening criteria show more variance (Table S1). The compression of values for JSD_cf_ around unity means that all scaffolds perform similarly. For this criterion, the ideal worst entry is still taken to be 0 (although no scaffold performed that poorly), rather than the lowest value followed by a rescaling to 0. This procedure thus prevents insignificant discrepancies between scaffolds from being amplified. Similarly, the values of 𝒟_*c*|*f*_ are all either essentially zero or very small, so we take the ideal worst entry to be 1 and the ideal best to be 0, to avoid ranking on statistically-insignificant differences. A server where the SMAA-TOPSIS algorithm may be run to rank the users’ own data can be found at http://bjork.phas.ubc.ca. The server can select and rank candidate leads from multiple competing screening tests that would otherwise make the selection of leads a nontrivial maximum-likelihood ranking problem. (*27*)

### 3.9 Cyclic peptide synthesis

Cyclic peptides were constructed by solid-phase synthesis (CPC Scientific), using headto-tail macrocyclization. Purity was confirmed by reversed-phase high-performance liquid chromatography (RP-HPLC) as *>* 95%. Molecular weight was confirmed by electrospray ionization mass spectrometry (ESI-MS). Solubility in water was confirmed by clear solution/absence of precipitate at 1 mg/mL. Peptides were conjugated to BSA through a cysteine side-chain.

### 3.10 Seeded aggregation experiments

The seeding activity of the conformational peptide epitope glycindel scaffolds was tested in a thioflavin T (ThT) aggregation assay, by measuring the fibrillogenic aggregation of *α*synuclein monomers with and without the addition of BSA-conjugated cyclic peptide over time. *α*-synuclein protein monomer was purchased from R-Peptide (product serial number: S-1001-1). The initial *α*-synuclein monomer concentration started from 100*µ*M, and the system was either seeded with 100nM BSA-conjugated cyclic peptides or unseeded as a control. A BSA-only control was not performed, however, previous studies of anti-oligomer antibody binding by surface plasmon resonance have used BSA reference surfaces as negative controls (*23, 24*), and BSA has also been shown in previous studies to inhibit (rather than enhance) the aggregation of transthyretin and A*β*_1*−*40_ (*100, 101*) The cyclic peptides introduced as additional seeds in this assay are listed in Table S2 (with rankings given in Table S1). They are P1 ((1,3)EKTK), P2 ((2,3)TKEQ), P21 ((1,4)TKEQ), P37 ((2,2)KTKE),

P45 ((4,2)KTKE) and P48 ((3,4)TKEQ). The system contains 25*µ*M Thioflavin T, 20mM Tris-HCl, and 100mM NaCl dissolved in PBS pH 7.4. The reaction volume is 120*µ*l. Incubation temperature is 37^*°*^C. The system was shaken for 20 minutes every 30 minutes prior to each reading. Fluorescence readout was measured by excitation at 440nm and emission at 486nm every half hour. Two independent experimental seeding assays were performed. Conditions in both assays were the same except the plate sealer, which resulted in more evaporation from samples in run 1, and lower signal for that run. Experimental data is provided in the Supporting Information.

The relative concentrations of aggregates were normalized by the concentration of the p2 peptide and fitted using the AmyloFit server (*43*). The model was chosen to be “Fragmentation Dominated, unseeded”, which was chosen as a minimal model that is sufficient to produce reasonable fitting (Fig. S3). The “unseeded” option here refers to the lack of any seeding by preformed fibril, and ensures that at the initial time, there is no fibril fragmentation. Since primary nucleation could be affected by the cyclic peptide seed, the nucleation rate (*k*_*n*_) was taken as the only parameter that varied across peptides for fitting the model. On the other hand, the fragmentation rate (*k*_2_) was fitted globally, and the critical nucleus size for primary nucleation (*n*_*c*_), and the number of monomer threshold to trigger secondary nucleation (*n*_2_) were set to the suggested default constant value of 2.

For comparing embedding depth of fibrils in cyclic peptide ensembles in the seeding assay, *α*-synuclein fibril structures deposited on RCSB PDB were obtained by searching the keyword “alpha synuclein fibril” and the sequence motif “EKTKEQ”; This resulted in 32 PDB fibril structures.

## 4. Conclusion

We proposed a virtual screening method that predicts cyclic peptide immunogen candidates with a desired conformation ensemble that has the potential to raise oligomer-selective antibodies. This screening method is applied to a pool of 48 cyclic peptide candidates, which scaffold the *α*-synuclein epitope in a restricted sequence space using flanked glycines and a cysteine for immunogen conjugation (“glycindel” scaffolds). The aim was to find cyclic peptide scaffolds that resemble the conformation of the epitope in the context of an oligomer. These cyclic peptides can then be used as immunogens to raise oligomer-selective antibodies. Other scaffolding and design strategies may be explored as well, to which the screening and ranking method developed here may be applied. In ongoing work, several highly-ranked cyclic peptides from this study have been used in active immunizations for the purpose of raising such oligomer-selective antibodies.

## Supporting information

Seeded aggregation ThT fluorescence data

## 5 Author Information

### 5.1 Author Contributions

Conceptualization, S.S.P.; Methodology, S.H., X.P., A.A. and S.S.P.; Software, X.P., S.H., and A.A.; Formal analysis, S.H., S.S.P., X.P., and A.A.; Experimental Data, A.Y.R.; Experimental Data Analysis: S.P. and S.H.; Resources, S.S.P and N.R.C.; Writing—original draft preparation, S.H., A.A., and S.S.P.; Writing—review and editing, S.S.P.; Visualization, S.H., X.P., A.A., and S.S.P.; Supervision, S.S.P.; Project administration, S.S.P.; Funding acquisition, S.S.P. We would like to also acknowledge a suggestion of Dr. Eugene Williams involving the seeding of aggregation by cyclic peptides.

### 5.3 Conflicts of Interest

S.S.P. was Chief Physics Officer of ProMIS Neurosciences until October 2020. N.R.C. is Chief Scientific Officer of ProMIS Neurosciences. S.S.P., N.R.C., and X.P. are co-inventors on international patent application PCT/CA2019/051434 (Publication WO/2020/073121, applicant being University of British Columbia). The patent application describes immunogens and epitopes in *α*-synuclein, antibodies to these epitopes, and methods of their making as well as their use. Patent applications owned by the University of British Columbia are licensed to ProMIS Neurosciences. The work presented was financially supported in part by ProMIS Neurosciences. S.H., A.A., X.P., A.Yu.R., N.R.C. and S.S.P. have received consultation compensation from ProMIS Neurosciences.

#### 6 Abbreviations

A*β*: Amyloid-*β*
AD: Alzheimer ‘s disease
ALS: Amyotropic lateral sclerosis
BSA: Bovine serum albumin
CC: Collective coordinate
CG: Coarse grained
Cryo-EM: Cryo-electron microscopy
CTE: Chronic traumatic encephalopathy
𝒟_*KL*_: Kullback–Leibler divergence
FRET: Förster resonance energy transfer
JSD: Jensen Shannon Divergence
KDE: Kernel density estimation
KLH: Keyhole limpet hemocyanin
MD: Molecular dynamics
MDS: Multidimensional scaling
NMR: Nuclear magnetic resonance
PD: Parkinson’s disease
PDB: Protein data bank
RMSD: Root mean squared deviation
RMSF: Root mean squared fluctuations
SASA: Solvent accessible surface area
SOD1: Superoxide dismutase 1
SPE: Stochastic proximity embedding
ThT: Thioflavin T

## 7 Associated Content

### Supporting Information

The Supporting Information is available free of charge at (URL).

1. Ranking criteria and rankings for *α*-synuclein epitope scaffolds; Convergence of JSD and embedding depth; Off-pathway targets for each epitope; Seeding aggregation experiment; experimental data and analysis (PDF).
2. Seeded aggregation ThT fluorescence data (XLSX).
3. A github repository link to the code required to perform head-to-tail cyclization of peptide sequences of interest is here: https://github.com/PlotkinLab/Backbone-linkage-of-cyclic-peptides

