## Supporting Info for "Optimizing epitope conformational ensembles using *α*-synuclein cyclic peptide “glycindel” scaffolds: A customized immunogen method for generating oligomer-selective antibodies for Parkinson’s disease"

§*Current address: Center for Quantum Technology Research, School of Physics, Beijing Institute of Technology, Beijing, China 100081*

E-mail:

### Table of contents

Tables

1. S1—Ranking criteria and rankings for $\alpha$-synuclein epitope scaffolds

2. S2—Embedding depth of PDB fibrils in cyclic peptide ensembles $\mathcal{D}_{f|c}$

Figures

1. S1—Convergence of dimensional reduction

2. S2—Off-pathway targets found in the RCSB

3. S3—Aggregation fluorescence data were fitted by four different models

4. S4—Scatter plot of the average embedding depth $\langle \mathcal{D}_{f|c} \rangle$ *vs.* the scaled nucleation rate

#### S.1 The ranking criteria and rankings for $\alpha$-synuclein epitope scaffolds

Table S1: Ranking criteria and rankings for $\alpha$-synuclein epitope scaffolds. The abbreviations $\text{JSD}_{cf}$, $\text{JSD}_{cm}$, $\text{JSD}_{cs}$, $\mathcal{D}_{c|m}$, and $\mathcal{D}_{c|s}$ correspond to $\text{JSD}_{\text{cyclic-fibril}}$, $\text{JSD}_{\text{cyclic-monomer}}$, $\text{JSD}_{\text{cyclic-stress}}$, $\mathcal{D}_{\text{cyclic-in-monomer}}$, and $\mathcal{D}_{\text{cyclic-in-stress}}$, respectively. OP corresponds to off-pathway targeting.

| Rank | scaffold | scaffold symbol | $\mathbf{JSD_{cf}}$ | $\mathbf{JSD_{cm}}$ | $\mathbf{JSD_{cs}}$ | $\mathcal{D}_{\mathbf{c}\vert\mathbf{f}}(\times 10^{-5})$ | $\mathcal{D}_{\mathbf{c}\vert\mathbf{m}}$ | $\mathcal{D}_{\mathbf{c}\vert\mathbf{s}}$ | OP |
|---|---|---|---|---|---|---|---|---|---|
| 0 | $I_{best}$ | $I_{best}$ | 1.00000 | 1.000 | 0.000 | 0 | 0.000 | 1.000 | 1.000 |
| 1 | CGEKTKGGG | (1,3)EKTK | 0.99979 | 0.897 | 0.751 | 0 | 0.052 | 0.107 | 1.000 |
| 2 | CGGEKTKGGG | (2,3)EKTK | 0.99978 | 0.849 | 0.765 | 0 | 0.093 | 0.147 | 0.979 |
| 3 | CGGGEKTKGG | (3,2)EKTK | 0.99979 | 0.850 | 0.768 | 0 | 0.104 | 0.124 | 0.959 |
| 4 | CGEKTKGG | (1,2)EKTK | 0.99979 | 0.980 | 0.916 | 0 | 0.009 | 0.058 | 0.986 |
| 5 | CGGTKEQGGG | (2,3)TKEQ | 0.99833 | 0.869 | 0.740 | 12 | 0.128 | 0.177 | 0.893 |
| 6 | CGGEKTKGGGG | (2,4)EKTK | 0.99978 | 0.822 | 0.700 | 0 | 0.094 | 0.160 | 0.892 |
| 7 | CGGGEKTKGGGG | (3,4)EKTK | 0.99955 | 0.686 | 0.675 | 0 | 0.235 | 0.225 | 0.971 |
| 8 | CGEKTKG | (1,1)EKTK | 0.99979 | 0.961 | 0.951 | 0 | 0.011 | 0.019 | 1.000 |
| 9 | CGGGGEKTKGG | (4,2)EKTK | 0.99977 | 0.804 | 0.786 | 0 | 0.140 | 0.135 | 0.952 |
| 10 | CGGEKTKGG | (2,2)EKTK | 0.99967 | 0.886 | 0.875 | 0 | 0.082 | 0.061 | 1.000 |
| 11 | CGGTKEQGGGG | (2,4)TKEQ | 0.99974 | 0.905 | 0.871 | 0 | 0.144 | 0.084 | 1.000 |
| 12 | CGEKTKGGGG | (1,4)EKTK | 0.99974 | 0.889 | 0.917 | 0 | 0.070 | 0.061 | 1.000 |
| 13 | CGGTKEQGG | (2,2)TKEQ | 0.99979 | 0.975 | 0.976 | 0 | 0.058 | 0.038 | 1.000 |
| 14 | CGGGEKTKGGG | (3,3)EKTK | 0.99979 | 0.737 | 0.742 | 0 | 0.170 | 0.142 | 0.958 |
| 15 | CGGKTKEG | (2,1)KTKE | 0.99979 | 0.996 | 0.999 | 0 | 0.010 | 0.000 | 1.000 |
| 16 | CGGGTKEQGG | (3,2)TKEQ | 0.99951 | 0.904 | 0.903 | 0 | 0.122 | 0.049 | 0.995 |
| 17 | CGGGTKEQGGG | (3,3)TKEQ | 0.99974 | 0.911 | 0.877 | 0 | 0.193 | 0.076 | 0.991 |
| 18 | CGGEKTKG | (2,1)EKTK | 0.99979 | 0.948 | 0.962 | 0 | 0.058 | 0.022 | 0.987 |
| 19 | CGGTKEQG | (2,1)TKEQ | 0.99979 | 0.995 | 1.000 | 0 | 0.015 | 0.001 | 0.991 |
| 20 | CGKTKEGG | (1,2)KTKE | 0.99979 | 0.976 | 0.998 | 0 | 0.018 | 0.001 | 1.000 |
| 21 | CGTKEQGGGG | (1,4)TKEQ | 0.99979 | 0.972 | 0.943 | 0 | 0.037 | 0.044 | 0.917 |
| 22 | CGKTKEG | (1,1)KTKE | 0.99978 | 0.966 | 0.999 | 0 | 0.030 | 0.000 | 1.000 |
| 23 | CGTKEQG | (1,1)TKEQ | 0.99979 | 0.967 | 0.995 | 0 | 0.084 | 0.011 | 1.000 |
| 24 | CGGGKTKEGG | (3,2)KTKE | 0.99979 | 0.954 | 0.995 | 0 | 0.034 | 0.002 | 0.984 |
| 25 | CGTKEQGG | (1,2)TKEQ | 0.99979 | 0.951 | 0.971 | 0 | 0.150 | 0.019 | 1.000 |
| 26 | CGGGGEKTKGGG | (4,3)EKTK | 0.99973 | 0.723 | 0.661 | 0 | 0.187 | 0.200 | 0.811 |
| 27 | CGGGEKTKG | (3,1)EKTK | 0.99979 | 0.964 | 0.991 | 0 | 0.043 | 0.004 | 0.910 |
| 28 | CGGGGTKEQGGG | (4,3)TKEQ | 0.99925 | 0.902 | 0.834 | 1 | 0.099 | 0.084 | 0.801 |
| 29 | CGGGGEKTKGGGG | (4,4)EKTK | 0.99931 | 0.509 | 0.649 | 0 | 0.452 | 0.331 | 1.000 |
| 30 | CGGGGTKEQG | (4,1)TKEQ | 0.99979 | 0.934 | 0.994 | 0 | 0.184 | 0.005 | 0.973 |
| 31 | CGKTKEGGG | (1,3)KTKE | 0.99956 | 0.917 | 0.997 | 7 | 0.142 | 0.000 | 0.944 |
| 32 | CGGGTKEQG | (3,1)TKEQ | 0.99979 | 0.925 | 0.991 | 0 | 0.100 | 0.007 | 0.896 |
| 33 | CGGGGKTKEG | (4,1)KTKE | 0.99969 | 0.816 | 0.962 | 0 | 0.196 | 0.013 | 0.987 |
| 34 | CGGGGEKTKG | (4,1)EKTK | 0.99979 | 0.879 | 0.919 | 0 | 0.091 | 0.035 | 0.834 |
| 35 | CGGGKTKEGGG | (3,3)KTKE | 0.99922 | 0.809 | 0.953 | 5 | 0.197 | 0.012 | 0.940 |
| 36 | CGGGGKTKEGGG | (4,3)KTKE | 0.99599 | 0.725 | 0.881 | 35 | 0.257 | 0.047 | 0.932 |
| 37 | CGGKTKEGG | (2,2)KTKE | 0.99873 | 0.859 | 0.942 | 13 | 0.349 | 0.056 | 0.933 |
| 38 | CGTKEQGGG | (1,3)TKEQ | 0.99978 | 0.922 | 0.936 | 0 | 0.070 | 0.054 | 0.732 |
| 39 | CGGGKTKEG | (3,1)KTKE | 0.99974 | 0.991 | 0.999 | 0 | 0.097 | 0.000 | 0.705 |
| 40 | CGGKTKEGGGG | (2,4)KTKE | 0.99791 | 0.708 | 0.947 | 6 | 0.312 | 0.017 | 0.884 |
| 41 | CGKTKEGGGG | (1,4)KTKE | 0.99929 | 0.757 | 0.969 | 0 | 0.239 | 0.004 | 0.777 |
| 42 | CGGGKTKEGGGG | (3,4)KTKE | 0.99958 | 0.706 | 0.901 | 0 | 0.548 | 0.089 | 0.963 |
| 43 | CGGGGTKEQGGGG | (4,4)TKEQ | 0.99861 | 0.840 | 0.726 | 2 | 0.142 | 0.144 | 0.519 |
| 44 | CGGKTKEGGG | (2,3)KTKE | 0.99966 | 0.822 | 0.976 | 0 | 0.091 | 0.004 | 0.591 |
| 45 | CGGGGKTKEGG | (4,2)KTKE | 0.99420 | 0.567 | 0.788 | 138 | 0.448 | 0.142 | 0.547 |
| 46 | CGGGGTKEQGG | (4,2)TKEQ | 0.99979 | 0.958 | 0.940 | 0 | 0.084 | 0.032 | 0.281 |
| 47 | CGGGGKTKEGGGG | (4,4)KTKE | 0.99445 | 0.586 | 0.829 | 90 | 0.267 | 0.045 | 0.415 |
| 48 | CGGGTKEQGGGG | (3,4)TKEQ | 0.99979 | 0.903 | 0.972 | 0 | 0.221 | 0.009 | 0.000 |
| 49 | $I_{worst}$ | $I_{worst}$ | 0.00000 | 0.000 | 1.000 | $10^5$ | 1.000 | 0.000 | 0.000 |

#### S.2 The convergence of JSD and embedding depth

Here, we show the convergence of the dimension reduction performance on the pairwise RMSD matrix that describes fibril, stressed fibril, monomer and (1,4)TKEQ ensemble. Stochastic Proximity Embedding (SPE) (*1*) is performed to calculate JSD in reduced dimension. The different degrees of convergence in 3-11 dimensions (See Supplemental Fig. S1a) motivated us to use a weighted average JSD across several dimensions, by weighting the JSD in each dimension by the inverse of the residual in that dimension. In practice this weights higher dimensions more than lower dimensions.

Multidimensional scaling (MDS) is performed to calculate embedding depth in reduced dimension. Since the residual quickly converged in dimensions of 3 or larger, (See Supplemental Fig. S1b), we simply perform the embedding depth measure in 3D.

The residual of the MDS and SPE are defined respectively as follows:

$$\text{Residual}_{MDS} = \frac{\sum_{i<j} \left(d_{ij} - r_{ij}\right)^2}{\sum_{i<j} r_{ij}^2} \tag{S1}$$

$$\text{Residual}_{SPE} = \sum_{i<j} \frac{\left(d_{ij} - r_{ij}\right)^2}{r_{ij}} / \sum_{i<j} r_{ij} \tag{S2}$$

In equations (S1) and (S2), index $i$ and $j$ correspond to two different rows in a pairwise RMSD matrix including the fibril ensemble, the stressed fibril ensemble, the monomer ensemble, and a given cyclic peptide scaffold ensemble. Before any dimensional reduction, the RMSD matrix is an $N \times N$ matrix, where $N$ is the sum of the number of configurations of the above four ensembles, and the indices $i$ and $j$ run from 1 to $N$. $d_{ij}$ and $r_{ij}$ are the Euclidean distance between $i^{th}$ and $j^{th}$ row of the RMSD matrix in reduced dimension and in the original dimension, respectively.

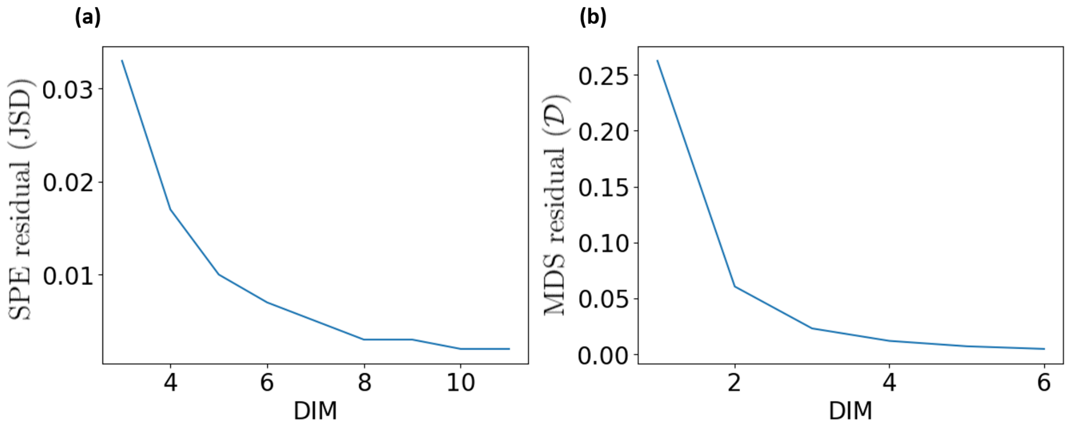


Figure S1: Convergence of dimensional reduction for (a) stochastic proximity embedding (SPE) for JSDs and (b) multidimensional scaling (MDS) for embedding depth. Both quantities are plotted vs. the dimension of the reduction. The scaffold used here for illustration is (1,4)TKEQ.

#### S.3 The off-pathway target for each epitope

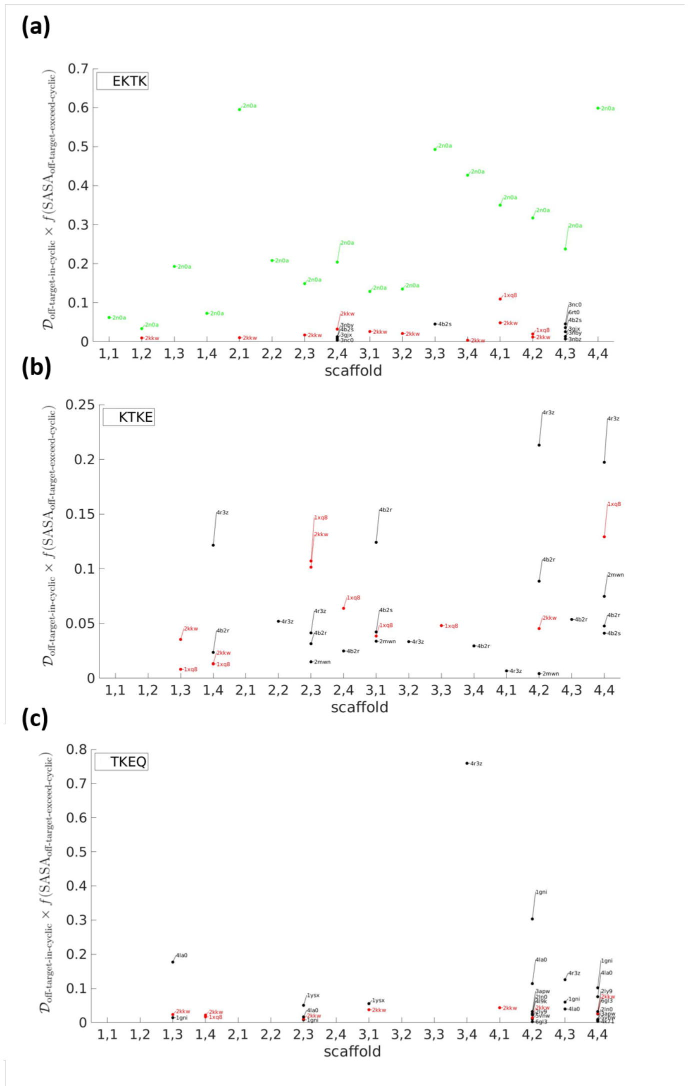


Figure S2: Off-pathway targets found in the RCSB, for each 4 amino acid epitope sequence and corresponding scaffold. E.g. 1,1 on the $x$-axis of the top panel is (1,1)EKTK. An off-pathway target is identified if both its $\mathcal{D}_{\text{off-target-in-cyclic}}$ and $f(\text{SASA}_{\text{off-target-exceed-cyclic}})$ are greater than 0.05, see e.g. Fig. 6c (main text). For such PDB entries, the value of each point on the $y$-axis is given by the product of $\mathcal{D}_{\text{off-target-in-cyclic}} \times f(\text{SASA}_{\text{off-target-exceed-cyclic}})$ for that off-pathway target, plotted here as a decimal value. The summation of values for points in the vertical column corresponding to each given scaffold is taken here to be the severity of the corresponding scaffold; This value is used to rank each scaffold. Results are presented for (a) EKTK (b) KTKE and (c) TKEQ epitopes. Entries colored green stand for fibrillar $\alpha$-synuclein (PDB 2N0A), while entries colored red are membrane-bound $\alpha$-synuclein monomers in the PDB, which typically contain $\alpha$-helical structure. The fibril $\alpha$-synuclein entries are specifically excluded from the off-pathway calculation because they have been treated already using JSD and embedding depth $\mathcal{D}$, while the structured monomers are retained in the off-pathway calculation.

### S.4 Seeding aggregation by cyclic peptides; experimental data and analysis

An Excel file of the seeded aggregation ThT fluorescence data used in this paper is included in the Supporting Information (“`seeding assays.xlsx`”).

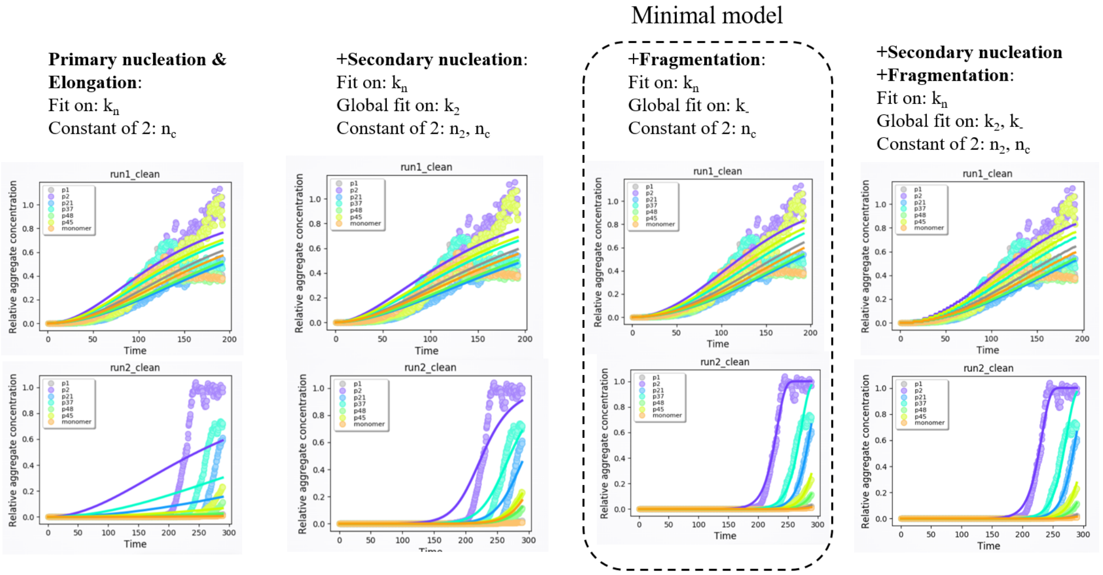


Figure S3: The two replicates (top row and bottom row) of aggregation fluorescence data were fitted by four different models. The results for each model are shown in each of the 4 columns. The left-most model only considers a primary nucleation ($k_n$) and elongation mechanism ($k_+$), which only have to be fitted separately in seeded fibril assay, and cannot produce good fit. The two models in the two middle columns consider the additional mechanisms of secondary nucleation ($k_2$) or fragmentation ($k_-$) respectively, with one additional parameter. The right-most model considers both secondary nucleation and fragmentation mechanisms. We found that fragmentation (in addition to primary nucleation and elongation) gave good quantitative fits to the data, while adding secondary nucleation did not. Moreover, adding secondary nucleation to a mechanism involving primary nucleation, elongation, and fragmentation did not further improve the fit to the data. As a result, a model that accounted for primary nucleation, elongation, and fragmentation was chosen as a minimal working model.

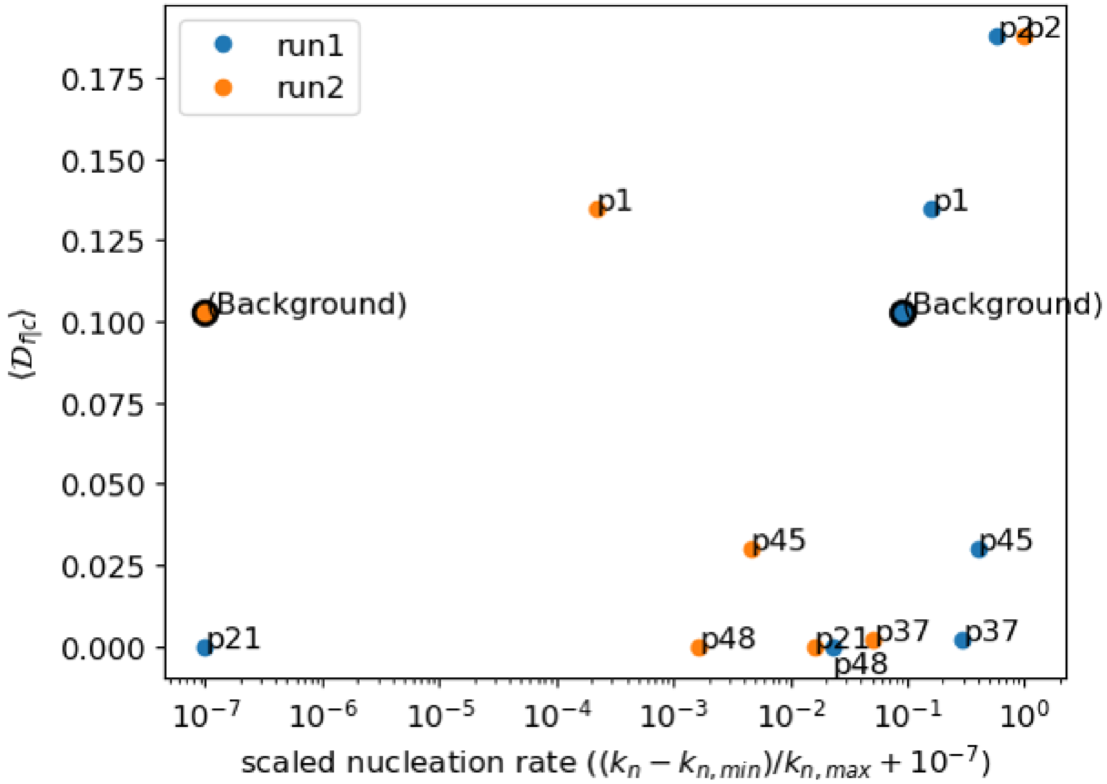


Figure S4: Scatter plot of the average embedding depth $\langle \mathcal{D}_{f|c} \rangle$ (Table S2), which measures the average similarity between PDB fibrils and each cyclic peptide, *vs.* the scaled nucleation rate from two separate seeding experiments. Nucleation rate is scaled by the minimum and maximum values in each separate experiment as shown on the axis label. The points labeled "(background)" correspond to no addition of cyclic peptide, and have an ordinate given by the average depth of the PDB fibrils in the monomer ensemble $\langle \mathcal{D}_{f|m} \rangle$.

Table S2: This table summarizes the embedding depth $\mathcal{D}_{f|c}$ of 32 PDB fibril structures of $\alpha$-synuclein in the 6 cyclic peptide ensembles examined in the seeded aggregation experiments, along with the embedding depth in the monomer ensemble ($\mathcal{D}_{f|m}$). The columns are ordered left to right by the nucleation rate fitted from the first replicate aggregation experiment (run 1). The fibril PDB names in bold have a monotonic decrease of $\mathcal{D}_{f|c}$ in the same order as the decreasing nucleation rate $k_n$, and underlined fibril names have 1 violation of this decreasing trend. 6RTB does not contain resolved TKEQ structure, so the depth related with TKEQ scaffolds cannot be calculated. Average $\mathcal{D}_{f|c}$ and nucleation rates $k_n$ are also given at the bottom of the table.

| Fibril PDB | P2 (2,3)TKEQ | P45 (4,2)KTKE | P37 (2,2)KTKE | P1 (1,3)EKTK | P48 (3,4)TKEQ | P21 (1,4)TKEQ | Background (monomer only) |
|---|---|---|---|---|---|---|---|
| 2N0A | 0.0167 | 0.0196 | 0 | 0 | 0 | 0 | 0.3986 |
| **6FLT** | 0.0721 | 0.0498 | 0.0005 | 0 | 0 | 0 | 0.0157 |
| 6L4S | 0.2448 | 0.014 | 0 | 0.8442 | 0 | 0 | 0.095 |
| 6OSM | 0.3552 | 0 | 0 | 0.0022 | 0 | 0 | 0.23 |
| 6RTB | 0 | 0 | 0 | 0 | - | - | 0.1534 |
| 6XYO | 0.1272 | 0.0109 | 0 | 0.1547 | 0 | 0 | 0.1272 |
| 7E0F | 0.1381 | 0.014 | 0 | 0.3804 | 0 | 0 | 0.0896 |
| 7NCH | 0.272 | 0.0203 | 0 | 0.3787 | 0 | 0 | 0.0453 |
| 6A6B | 0 | 0 | 0 | 0 | 0 | 0 | 0.0033 |
| 6H6B | 0.3527 | 0.0618 | 0 | 0.001 | 0 | 0 | 0.0256 |
| 6LRQ | 0 | 0 | 0 | 0 | 0 | 0 | 0.0075 |
| 6PEO | 0 | 0 | 0 | 0 | 0 | 0 | 0.0472 |
| 6SST | 0.1336 | 0.0185 | 0 | 0.216 | 0 | 0 | 0.1204 |
| 6XYP | 0.1288 | 0.0667 | 0.0031 | 0.2513 | 0 | 0 | 0.177 |
| 7LC9 | 0.0776 | 0 | 0 | 0.0195 | 0 | 0 | 0.2953 |
| 7NCI | 0.2283 | 0.0222 | 0.0002 | 0.3062 | 0 | 0 | 0.0493 |
| **6CU7** | 0.0648 | 0 | 0 | 0 | 0 | 0 | 0.1176 |
| 6L1T | 0.6877 | 0.0101 | 0.0037 | 0.2842 | 0 | 0 | 0.0361 |
| **6OSJ** | 0.033 | 0.0173 | 0 | 0 | 0 | 0 | 0.0664 |
| 6PES | 0 | 0 | 0 | 0 | 0 | 0 | 0.0155 |
| 6SSX | 0.0971 | 0.0167 | 0 | 0.0826 | 0 | 0 | 0.1924 |
| 6XYQ | 0.1295 | 0.0523 | 0 | 0.1377 | 0 | 0 | 0.1509 |
| 7NCA | 0.4127 | 0.0068 | 0.0003 | 0.0058 | 0 | 0 | 0.0649 |
| 7NCJ | 0.2832 | 0.0173 | 0.0004 | 0.2044 | 0 | 0 | 0.0786 |
| 6CU8 | 0.2017 | 0.2051 | 0.0086 | 0.0742 | 0 | 0 | 0.1634 |
| 6L1U | 0.4611 | 0.0001 | 0.017 | 0.1473 | 0 | 0 | 0.0417 |
| 6OSL | 0.4278 | 0.0533 | 0.0053 | 0.1264 | 0 | 0 | 0.0916 |
| 6RT0 | 0.0974 | 0.0065 | 0 | 0.1035 | 0 | 0 | 0.144 |
| 6UFR | 0.1423 | 0.0992 | 0 | 0.2382 | 0 | 0 | 0.0495 |
| 7C1D | 0.287 | 0.0086 | 0 | 0.6844 | 0 | 0 | 0.0355 |
| 7NCG | 0.3265 | 0.0937 | 0 | 0.0973 | 0 | 0 | 0.1011 |
| 7NCK | 0 | 0.0591 | 0.0218 | 0 | 0 | 0 | 0.0653 |
| average $\mathcal{D}_{f|c}$ | 0.1812 | 0.0295 | 0.0019 | 0.1481 | 0 | 0 | 0.103 |
| $k_n$ (run 1) | $4.81\times10^7$ | $3.92\times10^7$ | $3.44\times10^7$ | $2.79\times10^7$ | $2.12\times10^7$ | $1.58\times10^7$ | $2.44\times10^7$ |
| $k_n$ (run 2) | $2.28\times10^2$ | $1.09\times10^0$ | $1.17\times10^1$ | $1.01\times10^{-1}$ | $4.30\times10^{-1}$ | $3.70\times10^0$ | $5.07\times10^{-2}$ |
